# Differences between barley and maize revealed in limitations of photosystems I and II under high temperature and low air humidity

**DOI:** 10.1101/2024.05.16.594504

**Authors:** Eugene A. Lysenko

**Affiliations:** Institute of Plant Physiology, Russian Academy of Sciences; ul. Botanicheskaya 35, 127276, Moscow, Russia

**Author notes:** Corresponding Author:* Eugene A. Lysenko, Institute of Plant Physiology RAS, ul. Botanicheskaya 35, 127276, Moscow, Russia. . (E-mail address published in previous articles when EAL was the corresponding author.).

**Keywords:** maize, barley, chlorophyll fluorescence, P_700_ light absorption, limitations of photosystems, non-photochemical quenching, heat stress, air humidity.

## Abstract

Non-photochemical quenching and limitations of the photosystem I and photosystem II activities were studied in C_3_-plant barley and C_4_-plant maize. Plants were undergone to prolonged heat stress under high and low air humidity. Both species decreased non-photochemical quenching under tolerated heat stress (37-42°C), while increased it under nearly lethal heat stress (46°C). Usually, limitation at the acceptor side of the photosystem I was minor, while at 46°C it appeared major limiting factor. A similar decrease of photosystem II activity at 46°C by lower air humidity was achieved through different mechanisms. In barley, photosystem II downregulated by the increase of non-photochemical quenching. In maize, photosystem II downregulated by the increase of limitation at the acceptor side. Analysis of transients also revealed differences between species. One second after a light induction, limitations flashes at the acceptor sides of both photosystems. Elevating temperature reduced the size of these flashes; acceptor-side limitations of both photosystems decreased proportional to each other. In maize, the decrease was simple: the size of flashes slightly decreased at 37°C and more reduced at 42-46°C. In barley, the decrease had complex pattern: the size of flashes greatly reduced at 37°C and gradually returned to the control level under the higher temperatures. Around the photosystem II, the flash was quenched by a later burst of non-photochemical quenching. In barley, the transient peaks of acceptor-side limitation and non-photochemical quenching were very similar at any temperature. This was not observed in maize. The ratios between limitations qC/Y(NA) and qC/Y(ND) were studied.

**Highlights:**

- Light 1s induces instant limitations at acceptor-sides of PSII (qC) and PSI (Y(NA))
- Temperature reduced these flashes of qC and Y(NA) proportional to each other
- The pattern of reduction was different in barley and maize
- Flash of qC was quenched by proportional flash of qN in barley but not in maize
- Stationary PSII activity was decreased by different mechanisms in barley and maize

## Introduction

Plants with C_4_-photosynthesis have CO_2_-concentrating mechanism that had arisen multiple times in evolution of Angiosperms (Sage et al 2012). The largest portion of C_4_-species belongs to PACMAD clade of Poaceae family; C_4_-species represents 42% of all species in this large family (Togawa-Urakoshi and Ueno 2022). Among dicots, most C_4_-species belongs to Chenopodiaceae family (Edwards and Voznesenskaya 2011). Few algal lines also acquired C_4_-photosynthesis (Langdale 2011).

Specific features of plant C_4_-photosynthesis are Kranz anatomy (Haberlandt 1884) and Hatch-Slack cycle (Hatch and Slack 1966). Variations of Kranz anatomy are numerous, including its absence and single-cell C_4_-photosynthesis (Edwards and Voznesenskaya 2011). Biochemical variations of Hatch-Slack cycle are divided to three major groups depending on a major enzyme that releases primary fixed CO_2_ to Calvin-Benson cycle. There are three decarboxylases: chloroplastic NADP-dependent malic enzyme (EC 1.1.1.40; NADP-ME type), mitochondrial NAD-dependent malic enzyme (EC 1.1.1.39; NAD-ME type), and cytosolic phosphoenolpyruvate carboxykinase (EC 4.1.1.49; PEP-CK type) (Edwards and Voznesenskaya 2011; Langdale 2011). In NADP-ME type, bundle sheath (BS) cell chloroplasts have greatly reduced grana development; they are considered as mainly agranal (maize) or agranal (sorghum, sugarcane). In NADP-ME type, BS cells contains few, small mitochondria (Edwards and Voznesenskaya 2011). In NAD-ME type, BS cells contains bona fide granal chloroplasts and numerous, large mitochondria, while mesophyll (M) cells demonstrate some grana deficiency. In PEP-CK type, both M and BS cells have well-developed granal structure; BS cells have moderate abundance of small mitochondria (Edwards and Voznesenskaya 2011).

A theory predicts that C_3_-plants have more preferences under high CO_2_ level, low temperature, and low light conditions, while C_4_-plants have more advantages under low CO_2_ level, high temperature, and high light conditions (Johnson et al 2021). C_4_-plants have higher water- and nitrogen use efficiency than C_3_-plants (Makino et al 2003; Togawa-Urakoshi and Ueno 2022). Also, C_4_-plants perform net N mineralization of soil better than C_3_-plants (Reich et al 2018).

Up-to-date theory of C_4_-photosynthesis gives the fine knowledge of leaf anatomy, cell biology and CO_2_-metabolism. However, specific abilities of photosynthetic electron transport chain mostly remained out of focus. In this area, the contemporary knowledge is restricted to three points. First, grana-deficient chloroplasts have reduced content of the photosystem II (PSII). It was firmly demonstrated for BS chloroplasts in NADP-ME type: PSII subunits were absent (Kubicki et al 1996) or greatly reduced in numbers (Takabayashi et al 2005; Darie et al 2006). In NAD-ME type, M chloroplasts also have moderate grana-deficiency (Edwards and Voznesenskaya 2011); however, the decrease of PSII content was not obvious in these chloroplasts (Takabayashi et al 2005).

Second, grana-deficient chloroplasts have increased content of NDH dehydrogenase-like complex and, possibly, PGR5/PGRL1 complex. Both complexes function in cyclic electron flow around photosystem I (PSI); they represent two alternative pathways of electron transfer from ferredoxin (Fd) to plastoquinone (Yamori and Shikanai 2016). The content of NDH dehydrogenase-like complex increased in grana-deficient BS chloroplasts in NADP-ME type (Kubicki et al 1996; Takabayashi et al 2005; Darie et al 2006; Munekage et al 2010; Nakamura et al 2013) and in grana-deficient M chloroplasts in NAD-ME type (Takabayashi et al 2005). Among species of Flavaria genus, PGR5 and PGRL1 contents are higher in NADP-ME type C_4_-species comparing with intermediate C_3_-C_4_ and regular C_3_ species (Munekage et al 2010; Nakamura et al 2013). However, other dicotyledonous species and maize demonstrated no difference in PGR5 content among M chloroplasts of C_3_-species and BS and M chloroplasts of C_4_-species (Takabayashi et al 2005).

Third, maximal quantum yield of PSII Fv/Fm in C_3_-plants is higher than in C_4_-plants of NADP-ME type (Kalaji et al 2014). The method of pulse amplitude modulation (PAM) is fine tool for analysis of photosynthetic electron transport chain (Kalaji et al 2014). Analysis of chlorophyll (Chl) fluorescence revealed some differences in PSII functioning between NADP-ME and NAD-ME types of C_4_-plants (Takabayashi et al 2005). Analysis of P_700_-oxidation state showed larger impact of cyclic electron flow around PSI in C_4_-species comparing with C_3_-species (Nakamura et al 2013). Both studies analyzed fast (transient) reactions of PSI and PSII activities to changes of illumination.

Barley is classic model C_3_-species. Maize is classic model C_4_-species with a rare feature: its major NADP-ME pathway is accomplished with PEP-CK as a minor pathway (Walker et al 1997; Koteyeva et al 2015). Recently, we performed comparative studies of barley and maize under different temperatures and water contents in air. In this study, we demonstrated that lower air humidity inhibits both plant growth and photochemical activities of PSII and PSI; this influence was of critical importance under severe heat stress (HS) (Lysenko et al 2023). In this article, factors limiting the photochemical activities of PSII and PSI remained out of consideration because they should be mostly examined from the standpoint of species specificity.

The current study has completed the whole analysis. Young barley and maize plants were treated for 48 h under elevated temperatures (24/37/42/46°C) and contrast water contents in air - high humidity (HH) and low humidity (LH). Plants were measured with Dual-PAM-100 under a temperature of 48 h treatment. The current analyses focused on limitations of PSI at the donor-side (Y(ND)) and the acceptor-side (Y(NA)), limitation of PSII at the acceptor-side (qC), and non-photochemical quenching by PSII. The special attention paid to fast (transient) reactions of PSI and PSII to a light induction. The original experimental design and standpoint of analyses enabled describing many features of PSI and PSII functioning for the first time. Three features occurred to be of special interest. First, the hardest stress (46°C, LH) similarly reduced the steady-state level of PSII photochemical activity in both species (Lysenko et al 2023); however, in C_3_-plant barley it was achieved through an up-regulation of non-photochemical quenching (qN) while in C_4_-plant maize it was caused with an increase in acceptor-side limitation (qC). Second, a transient peak of PSII acceptor-side limitation (qC) was fast resolved with a subsequent transient peak of non-photochemical quenching (qN) at that amplitudes of the former and the later were proportional; this proportional quenching was observed in C_3_-plant barley but not in C_4_-plant maize. Third, transient peaks of acceptor-side limitations of PSII (qC) and PSI (Y(NA)) varied at wide interval remaining proportional to each other; this proportionality was observed in both species though the ratios were different in barley and maize. The comprehensive analysis of limitations and non-photochemical quenching is presented below.

## 2. Material and methods

The current work and the article (Lysenko et al., 2023) represents analysis of the single experiment. The earlier study has been focused on the plant growth and the photochemical activities of PSI and PSII (Lysenko et al., 2023); the current study focuses on limitations of PSI and PSII activities. The whole experiment has been described comprehensively in (Lysenko et al., 2023); this manuscript outlines the experiment briefly. For any details, readers are referred to (Lysenko et al., 2023).

### 2.1. Plant growth conditions

Maize (*Zea mays* L. cv. Luchistaya) and barley (*Hordeum vulgare* L. cv. Luch) seedlings were grown in phytotron chambers at 21°C (barley) or 25-26°C (maize), 180–200 μmol photons m^-2^ s^-1^, and a photoperiod of 16 h light/8 h dark under continuous aeration on modified Hoagland medium. Seven-day-old seedlings were transferred to thermostat chambers for 48 h; they were grown under continuous illumination at 60-80 μmol photons m^-2^ s^-1^ and temperatures 24°C, 37°C, 42°C, or 46°C. Each of the four temperature regimes was accomplished in parallel at two different levels of relative air humidity (HH and LH, Lysenko et al., 2023). Under the most severe conditions (46°C and LH), barley plants were dying fast after 51-52 h of treatment.

### 2.2. PAM analysis

The Chl *a* fluorescence and P_700_ light absorption were registered simultaneously with the Dual-PAM-100 (Walz, Germany). The largest fully developed leaf was used for the measurement: the first leaf of barley and second leaf of maize. The Chl *a* fluorescence was excited at 460 nm (9 μmol photons m^−2^ s^−1^; the measuring light). P_700_ is the reaction center chlorophyll of PSI; the level of oxidized P_700_ was measured as the difference in light absorption at 830 and 875 nm. The red light 635 nm was used as the actinic light (AL) and for the saturation pulses (SPs). In each variant, the thermal treatment (48 h), adaptation to the darkness (30 min), and all the measurements with Dual-PAM-100 were performed in thermostat chambers under the same temperature (24°C, 37°C, 42°C, or 46°C).

In the dark, the measuring light only was used for the determination of minimal (Fo) Chl fluorescence; the first SP was applied for the measurement of maximal (Fm) Chl fluorescence; the pre-illumination with far-red light (720 nm, Int. #10, ∼250 μmol photons m^-2^ s^-1^, Pfündel et al., 2013) followed with the second SP were employed for the determination of minimal (Po) and maximal (Pm) P_700_ absorption.

Next, the AL 70 μmol photons m^-2^ s^-1^ was induced for 7.5 min and induction curves (ICs) were recorded; SPs (4 mmol photons m^−2^ s^−1^, 500 ms) were induced every 40 s and followed by far-red light (720 nm, Int. #10, ∼250 μmol photons m^-2^ s^-1^, Pfündel et al., 2013) for 5 s. The maximum level of fluorescence in the light (Fm’) and maximal P_700_ change in the light (Pm’) were measured with SPs. The minimum level of fluorescence in the light (Fo’) was determined with the far-red illumination. The stationary Chl fluorescence (Fs) and P_700_ absorption (P) were measured just before SP (Pfündel et al., 2013). The minimal level of P_700_ light absorption (Po) was measured after cessation of far-red light both in the dark and light regimes (Klughammer and Schreiber, 1994, 2008).

After the end of IC measurement, the measurement of rapid light curve (RLC) was started immediately according to (Lysenko, 2021). In RLC, each step of AL lasted for 30 s; the set of AL intensities is shown in the figures. The other parameters were the same as in IC. The method RLC aspires to but does not reach a stationary level (White and Critchley 1999; Kalaji et al 2014); RLC data should be considered as quasi-stationary.

Limitations of the photochemical activity of PSI were calculated as follows: limitation at the donor-side Y(ND) = (P – Po)/(Pm – Po) and limitation at the acceptor-side Y(NA) = (Pm – Pm’)/(Pm – Po) (Klughammer and Schreiber, 1994, 2008). Limitation of the photochemical activity of PSII at the acceptor side (“closed PSII”) qC = (Fs – Fo’)/Fv (Lysenko et al., 2020). Non-photochemical quenching of light energy in PSII qN = (Fv – Fv’)/Fv (Schreiber et al., 1986; van Kooten and Snel, 1990) where Fv = Fm – Fo and Fv’ = Fm’ – Fo’. qN + X(II) + qC = 1 (Lysenko et al., 2020).

Calculation of the ratios qC/Y(ND) and qC/Y(NA) were performed according to (Lysenko and Kusnetsov 2024). Particular ratios with negative, zero, or extremely small denominators (/Y(ND) or /Y(NA)) were discriminated; the discrimination was solely restricted to the ratios qC/Y(ND) and qC/Y(NA). The general map of qC/Y(ND) and qC/Y(NA) data points discarded is represented in Suppl. Tables S1-S4.

### 2.3. Statistics

For each variant of *in vivo* treatment (48 h), biological experiments were repeated four-five times and more then 20 individual plants were analyzed. The most of Y(NA), Y(ND), qC, and qN curves represents an averages of 20 and more individual curves; the minimal number of individual curves was 23 (24°C), 19 (37°C), and 16 (42°C, 46°C).

The data were processed using the Excel (Microsoft) software. The significance of differences between mean values was verified using the two-tailed Student’s *t*-test.

## 3. Results

In the current study, each data set is visualized by two alternative modes to trace either the effect of temperature or the effect of air humidity; the effect of species specificity can be seen at both variants. For each data set, one way of visualization is shown in the article while alternative variant is given in the supplementary file.

### 3.1. Limitations of photosystems

The analysis of limitations is beneficial to begin from the endpoint of electron transfer chain. Limitation of PSI at the acceptor side Y(NA) was usually minor component restricting PSI activity (Fig. 1). There were two exceptions. First, switching AL on immediately induced flash of Y(NA). In control plants, illumination for 1 s increased Y(NA) from zero level to nearly 90% in barley and nearly 100% in maize. This burst was transient and 40 s later it mainly disappeared. Under the elevated temperatures, plants of both species acquired ability relaxing this flash of PSI limitation; however, barley and maize performed it differently. In maize, the maximal height of Y(NA) flash was slightly reduced at 37°C and more reduced at 42-46°C (Fig. 1E,G). In barley, Y(NA) flash was relaxed at 37°C greatly; the further increase of temperature gradually exhausted this ability and turned the size of Y(NA) flash back to the control level (Fig. 1A,C).

**Fig. 1.**
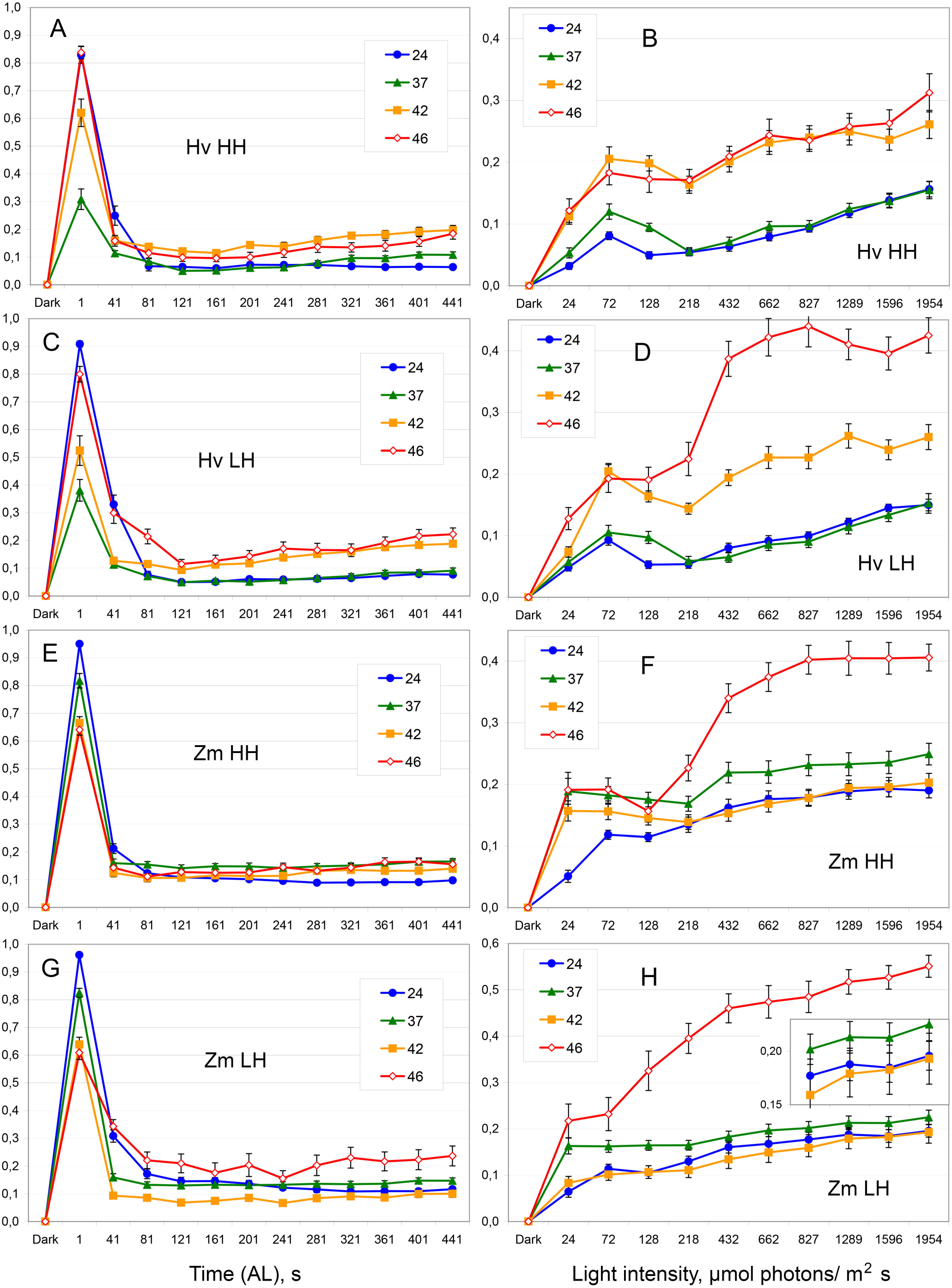
Limitation at the acceptor side of PSI (Y(NA)). A-D – barley (Hv); E-H – maize (Zm). A, C, E, G – IC (AL 70 μmol photons m^-2^ s^-1^); B, D, F, H – RLC. A-B, E-F – higher relative humidity of air (HH); C-D, G-H – lower relative humidity of air (LH). Filled blue circles – 24°C; filled green triangles – 37°C; filled orange squares – 42°C; open red diamonds – 46°C. The data are presented as means ± standard error (SE). The inset shows the corresponding small values with the higher resolution. The same data are presented in an alternative manner for analyzing the effect of air humidity (Suppl. Fig. S1).

Second, a large increase of Y(NA) was observed in plants grown at 46°C. In RLC, Y(NA) reached ∼40% at rather moderate AL intensity 432 μmol photons m^-2^ s^-1^ and higher (Fig. 1B,D,F,H). This increase was larger in maize and under LH conditions (Suppl. Fig. 1H); Y(NA) exceeded 50% in maize plants under LH conditions and at AL intensities >1 mmol photons m^-2^ s^-1^ (Fig. 1H, Suppl. Fig. 1H). In barley, a small increase of Y(NA) was also observed at 42°C (Fig. 1A-D).

Immediately (1 s) after AL induction, limitation of PSI at the donor side Y(ND) was relaxed nearly completely; then, Y(ND) increased in 1-2 min and remained at this level or more frequently decreased once again (Fig. 2A,C,E,G). In the first 40 s of illumination, Y(ND) was increasing similarly in both species (Suppl. Fig. S2A,C,E,G). In barley, Y(ND) grew until 80 or 120 s and reached the higher values; further, it was reduced even more comparing with maize (Suppl. Fig. S2A,C,E,G). In barley, elevated temperature slowed this transient increase; the peak of Y(ND) was shifted from 80 s (control) to 120 s (HS) (Fig. 2A,C,E,G).

**Fig. 2.**
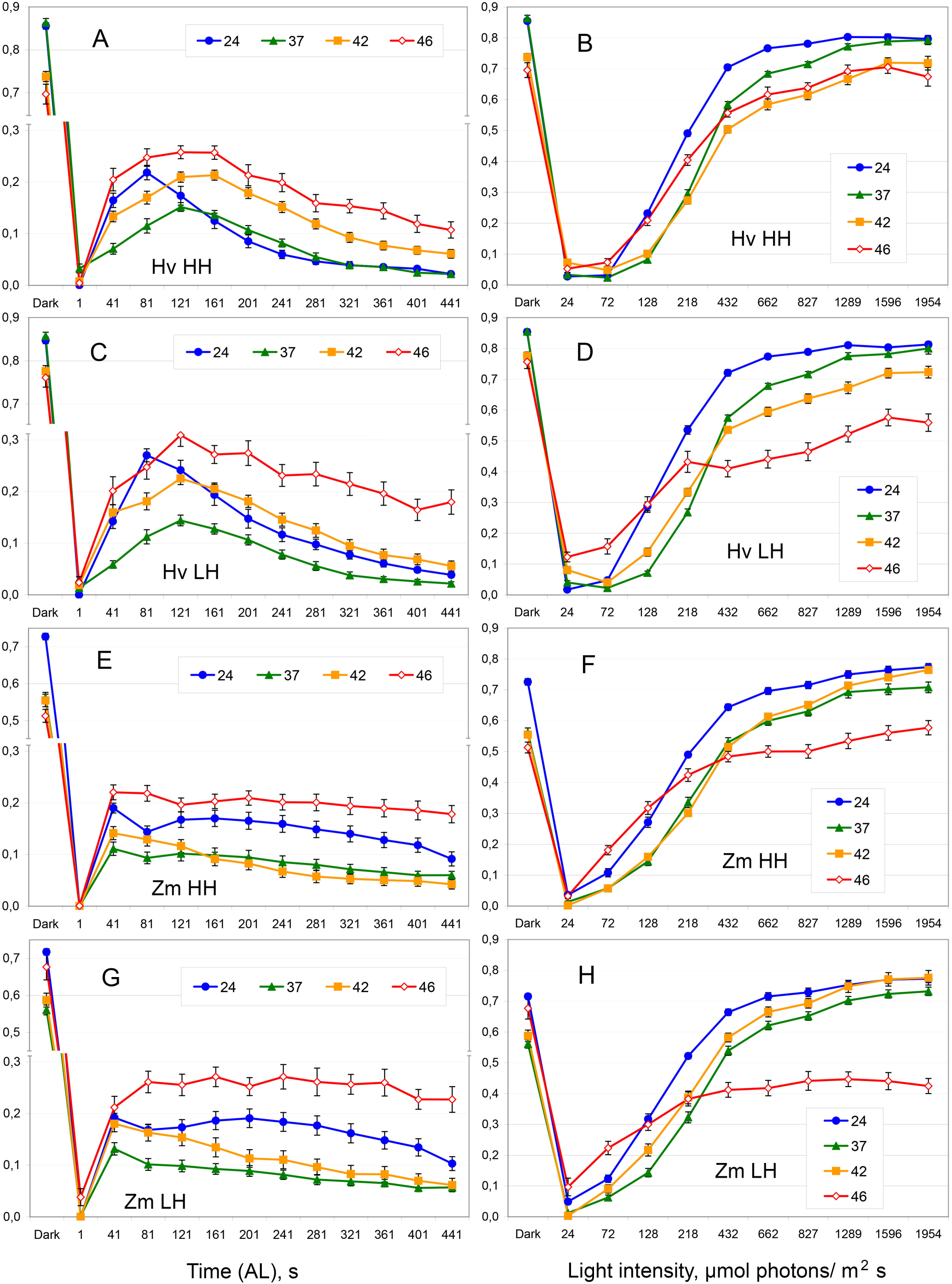
Limitation at the donor side of PSI (Y(ND)). A-D – barley; E-H – maize. A, C, E, G – IC; B, D, F, H – RLC. A-B, E-F – higher relative humidity of air; C-D, G-H – lower relative humidity of air. All designations are the same as in Fig. 1. The same data are presented in an alternative manner for analyzing the effect of air humidity (Suppl. Fig. S2).

Tolerated HS reduced Y(ND). At 37°C, Y(ND) always decreased (Fig. 2). At 42°C, Y(ND) mostly decreased (Fig. 2) and solely increased in IC of barley plants under HH conditions (Fig. 2A). Under nearly lethal HS (46°C), Y(ND) increased in IC at 72 μmol photons m^-2^ s^-1^ (Fig. 2A,C,E,G) and in RLC at low AL intensities (24-72 μmol photons m^-2^ s^-1^) (Fig. 2B,D,F,H). In RLC at AL 128-218 μmol photons m^-2^ s^-1^, Y(ND) was close to the control level; at the higher AL intensities, Y(ND) greatly decreased (Fig. 2B,D,F,H). Probably, the latter effect was caused with the great increase of Y(NA) under these conditions (Fig. 1B,D,F,H, Suppl. Fig. S1H). At 46°C, LH conditions enlarged the decrease of Y(ND) at AL intensities ≥ 432 μmol photons m^-2^ s^-1^ (Suppl. Fig. S2H); probably, it was also achieved through the great increase of Y(NA) (Suppl. Fig. S1H).

Limitations at the acceptor sides of PSII (qC) and PSI (Y(NA)) were rather similar. Both qC and Y(NA) showed a flash of limitation after AL induction and then usually demonstrated rather small values (Fig. 1, Fig. 3). In control plants, illumination for 1 s increased qC from zero level to nearly 60% in barley and nearly 80% in maize. Plants adapted to the elevated temperatures relaxed this flash of PSII limitation. Maize plants slightly reduced the peak at 37°C and more reduced the peak at 42-46°C (Fig. 3E,G). Barley plants greatly relaxed the flash of qC at 37°C; at the higher temperatures, this ability gradually exhausted and the size of qC flash turned back to the control level (Fig. 3A,C). In RLC, qC increased till AL intensities 432-662 μmol photons m^-2^ s^-1^ and then decreased at the higher AL intensities (Fig. 3B,D,F,H).

**Fig. 3.**
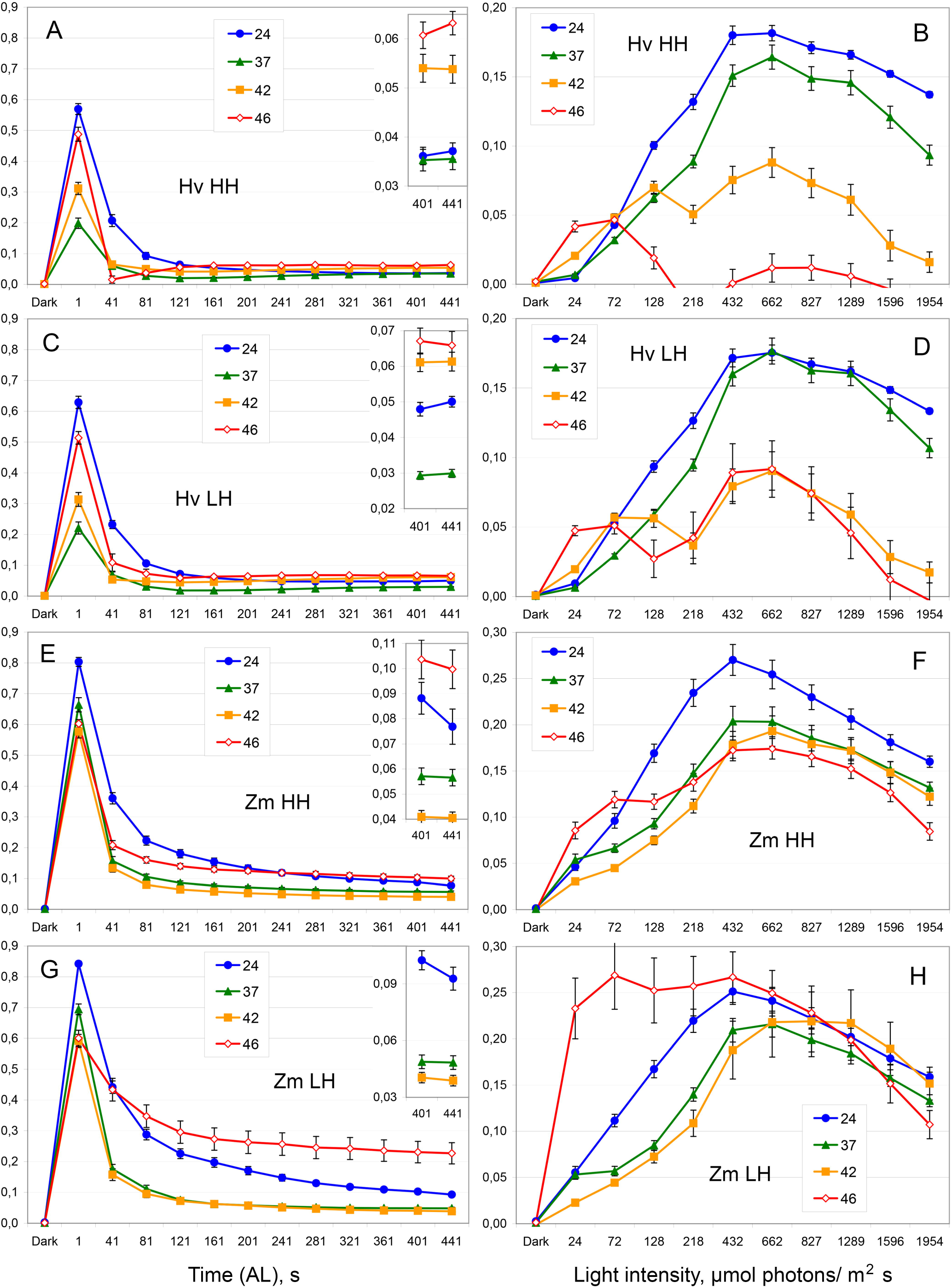
Limitation at the acceptor side of PSII (qC, “closed” PSII). A-D – barley; E-H – maize. A, C, E, G – IC; B, D, F, H – RLC. A-B, E-F – higher relative humidity of air; C-D, G-H – lower relative humidity of air. All designations are the same as in Fig. 1. The insets show the corresponding small values with the higher resolution. The same data are presented in an alternative manner for analyzing the effect of air humidity (Suppl. Fig. S3).

Maize plants, decreased qC at tolerated HS (37°C, 42°C) in both IC and RLC (Fig. 3E-H). Barley plants demonstrated no clear decrease of qC in IC at AL 72 μmol photons m^-2^ s^-1^ (Fig. 3A,C); the steady-state level was low in all the variants. In RLC, barley plants decreased qC at 42°C and 46°C; qC decrease was observed at AL ≥ 128 μmol photons m^-2^ s^-1^ (Fig. 3B,D).

The conditions of LH increased qC solely at 46°C (Suppl. Fig. 3). In maize, LH conditions unambiguously increased qC in both IC and RLC (significantly at AL 24-827 μmol photons m^-2^ s^-1^) (Suppl. Fig. 3G-H). In barley, LH conditions significantly increased qC only at AL 432-827 μmol photons m^-2^ s^-1^) (Suppl. Fig. 3H).

### 3.2. Non-photochemical quenching

In the beginning of IC, barley plants induced a peak of qN resembling the flash of qC in corresponding variants; however, the peak of qN was observed for 40 s later then the flash of qC (Fig. 4A,C). In RLC at 24 and 72 μmol photons m^-2^ s^-1^, a “shelf” of qN was observed in barley plants grown at 24°C and 37°C (Fig. 4B,D, Suppl. Fig. S4B,D); in all the other cases, an increase of AL intensity was followed with an increase of qN (Fig. 4, Suppl. Fig. S4). Light intensities influenced non-photochemical quenching in barley more than in maize; qN in barley was smaller under low AL and larger under high AL comparing with maize (Suppl. Fig. S4).

**Fig. 4.**
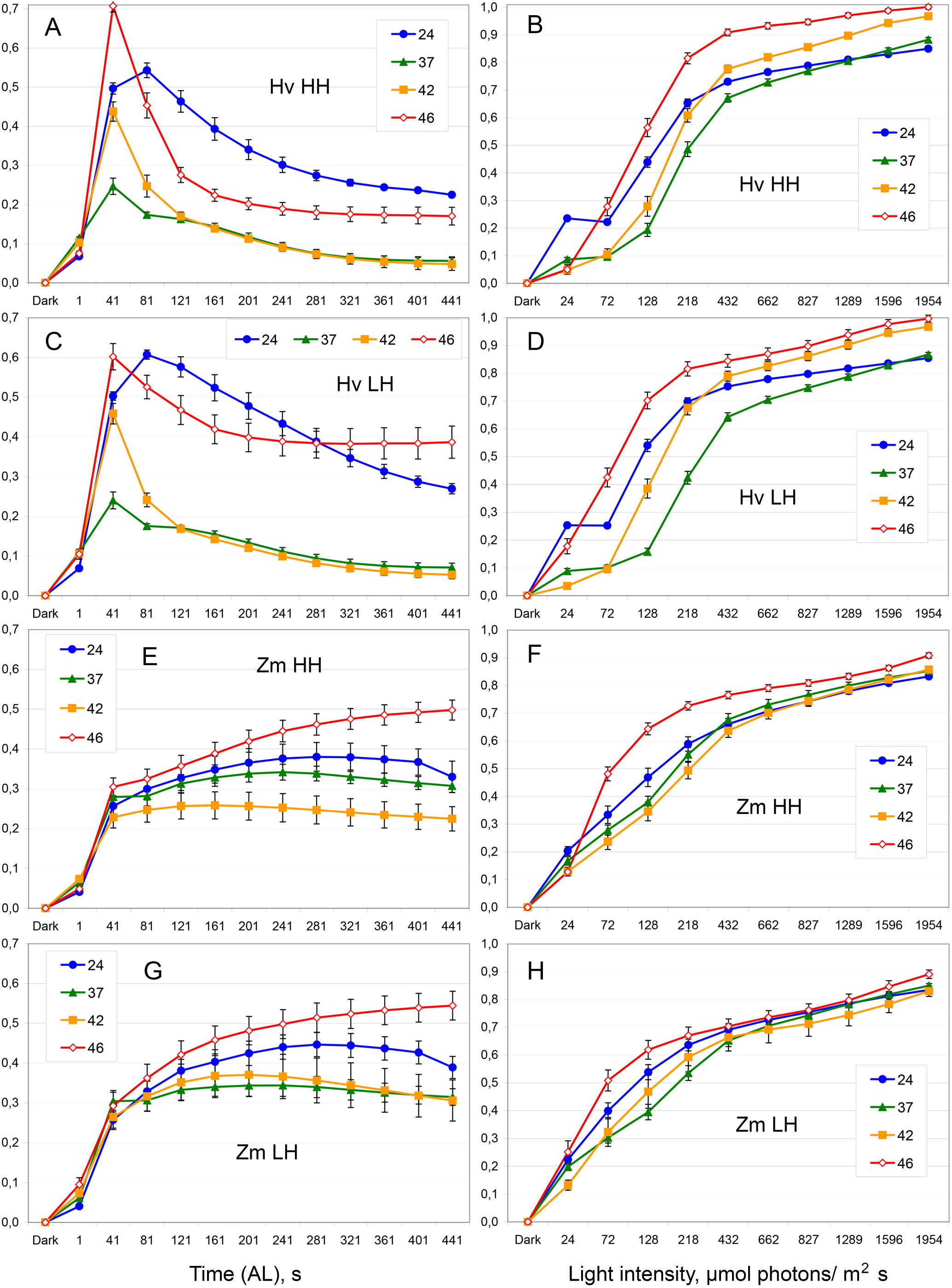
Non-photochemical quenching of PSII (qN). A-D – barley; E-H – maize. A, C, E, G – IC; B, D, F, H – RLC. A-B, E-F – higher relative humidity of air; C-D, G-H – lower relative humidity of air. All designations are the same as in Fig. 1. The same data are presented in an alternative manner for analyzing the effect of air humidity (Suppl. Fig. S4).

Both plant species demonstrated a general tendency decreasing non-photochemical quenching under tolerated HS (37-42 °C) and increasing qN under nearly lethal HS (46°C) and LH conditions (Fig. 4, Suppl. Fig. S4). Maize plants showed rather small changes of qN under different temperatures and air humidity (Fig. 4E-H, Suppl. Fig. S4) while barley plants manifested large diverse changes of non-photochemical quenching (Fig. 4A-D, Suppl. Fig. S4).

Under low AL intensities, LH conditions increased non-photochemical quenching in barley plants; a moderate increase was observed in the control and a rather large increase was found at 46°C. In IC, qN increase was obvious (Suppl. Fig. S4A,G). In RLC, qN increase in the control was small but significant at AL 24-218 μmol photons m^-2^ s^-1^ (Suppl. Fig. S4B); at 46°C, qN increase was rather large and significant at AL 24-128 μmol photons m^-2^ s^-1^ (Suppl. Fig. S4H, small decrease at 432-827 μmol photons m^-2^ s^-1^ also was significant).

### 3.3. Balance between limitations of PSI and PSII

Recently, we concluded that qC/Y(NA) ratio mostly reflects dynamics of qC (Lysenko and Kusnetsov 2024). In the current study, the dynamics of qC/Y(NA) ratio were distinct from the corresponding qC and Y(NA) dynamics (Fig. 5, Suppl. Fig. S5); solely, RLC of barley plants at 24°C and 37°C showed some similarity between qC/Y(NA) (Fig. 5B,D) and qC (Suppl. Fig. S3B,D) because Y(NA) in these variants fluctuated close to 0.1 (Suppl. Fig. S1B,D).

**Fig. 5.**
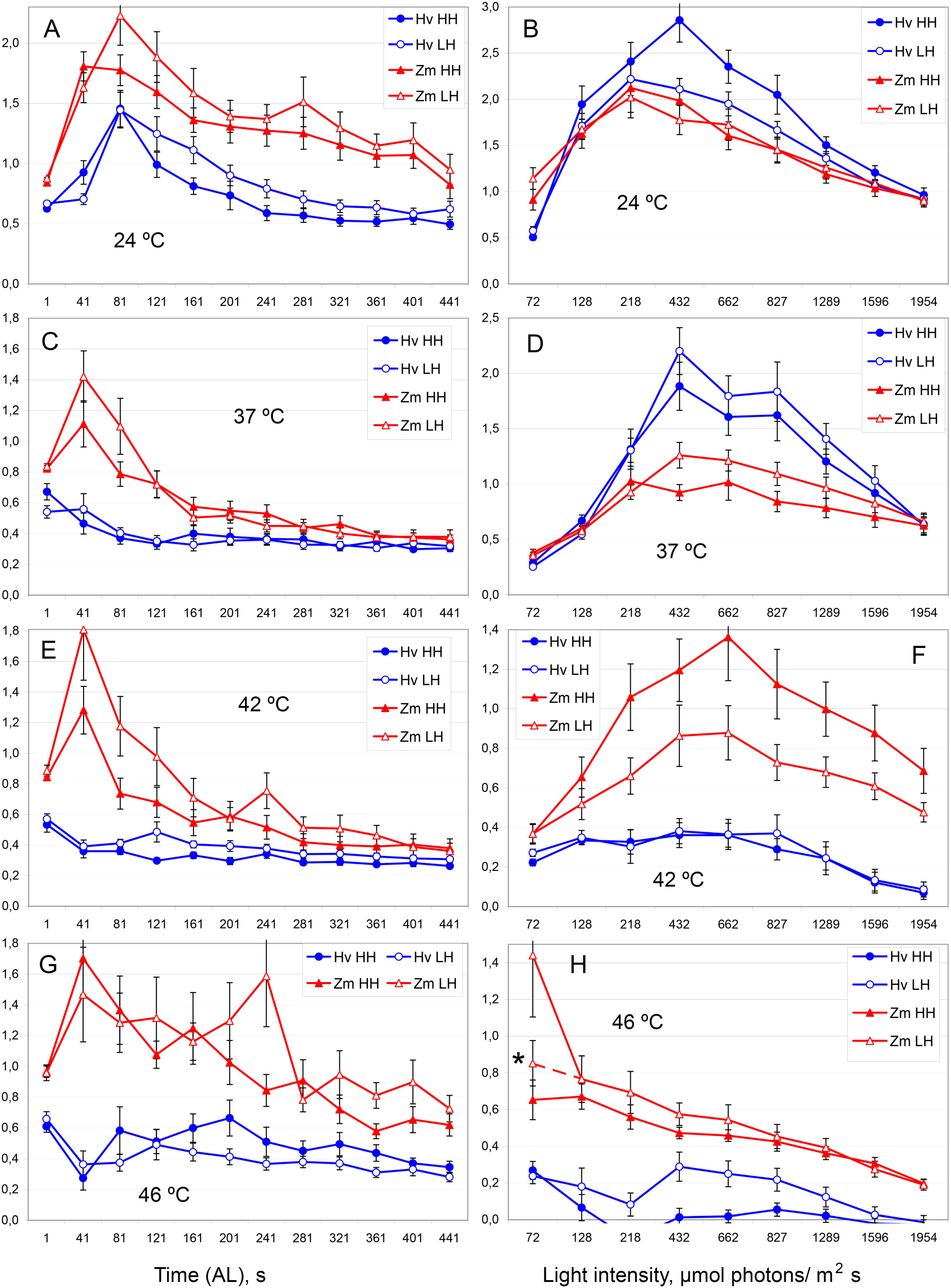
The ratio qC/Y(NA) showing balance between limitations at the acceptor sides of PSII (qC) and PSI (Y(NA)). A, C, E, G – IC (AL 70 μmol photons m^-2^ s^-1^); B, D, F, H – RLC. A, B – 24°C; C, D – 37°C; E, F – 42°C; G, H – 46°C. Blue lines and circles represent barley (Hv); red lines and triangles represent maize (Zm). * – variant of the corresponding point in curve without 3 drastically different but irremovable data points (as more credible; connected with dashed line). Filled symbols show HH conditions; open symbols show LH conditions. The data are presented as means ± SE. The same data are presented in an alternative manner for analyzing the temperature effect (Suppl. Fig. S5).

In both species, qC/Y(NA) ratios were tightly correlated immediately (1 s) after AL induction (Suppl. Fig. S5A,C,E,G); further, qC/Y(NA) changed diversely (Suppl. Fig. S5). The ratios qC/Y(NA) were both higher and lower than 1 (Fig. 5); hence, acceptor-side limitation of PSII can be larger or smaller than acceptor-side limitation of PSI. Generally, qC/Y(NA) ratio in maize was higher comparing with barley (Fig. 5). In RLC at 24°C and 37°C, qC/Y(NA) ratios were higher (or equal) in barley than in maize (Fig. 5B,D) while both qC and Y(NA) were larger in maize than in barley (Suppl. Fig. S1B,D, Suppl. Fig. S3B,D). In the beginning of IC, control plants showed peak of qC/Y(NA) at 81 s in barley and at 41 s or 81 s in maize (Fig. 5A); under HS, these peaks disappeared in barley and shifted completely to 41 s in maize (Fig. 5C,E,G). Elevating temperature decreased qC/Y(NA) ratio (Suppl. Fig. S5).

The ratio qC/Y(ND) enables comparing limitations between PSII and PSI. Similar to previous results (Lysenko and Kusnetsov 2024), qC/Y(ND) demonstrated complex pattern of changes and most values were ≤ 1; qC/Y(ND) > 1 were observed in maize mostly in the beginning of IC (Fig. 6). Generally, the ratios of qC/Y(ND) were higher in maize than in barley (Fig. 6). A short quasi-stationary fragments were observed in qC/Y(ND) dynamics; however, in the current study quasi-stationary fragments were less obvious (Fig. 6) than in our recent research (Lysenko and Kusnetsov 2024). This topic will be thoroughly considered in the Discussion chapter.

**Fig. 6.**
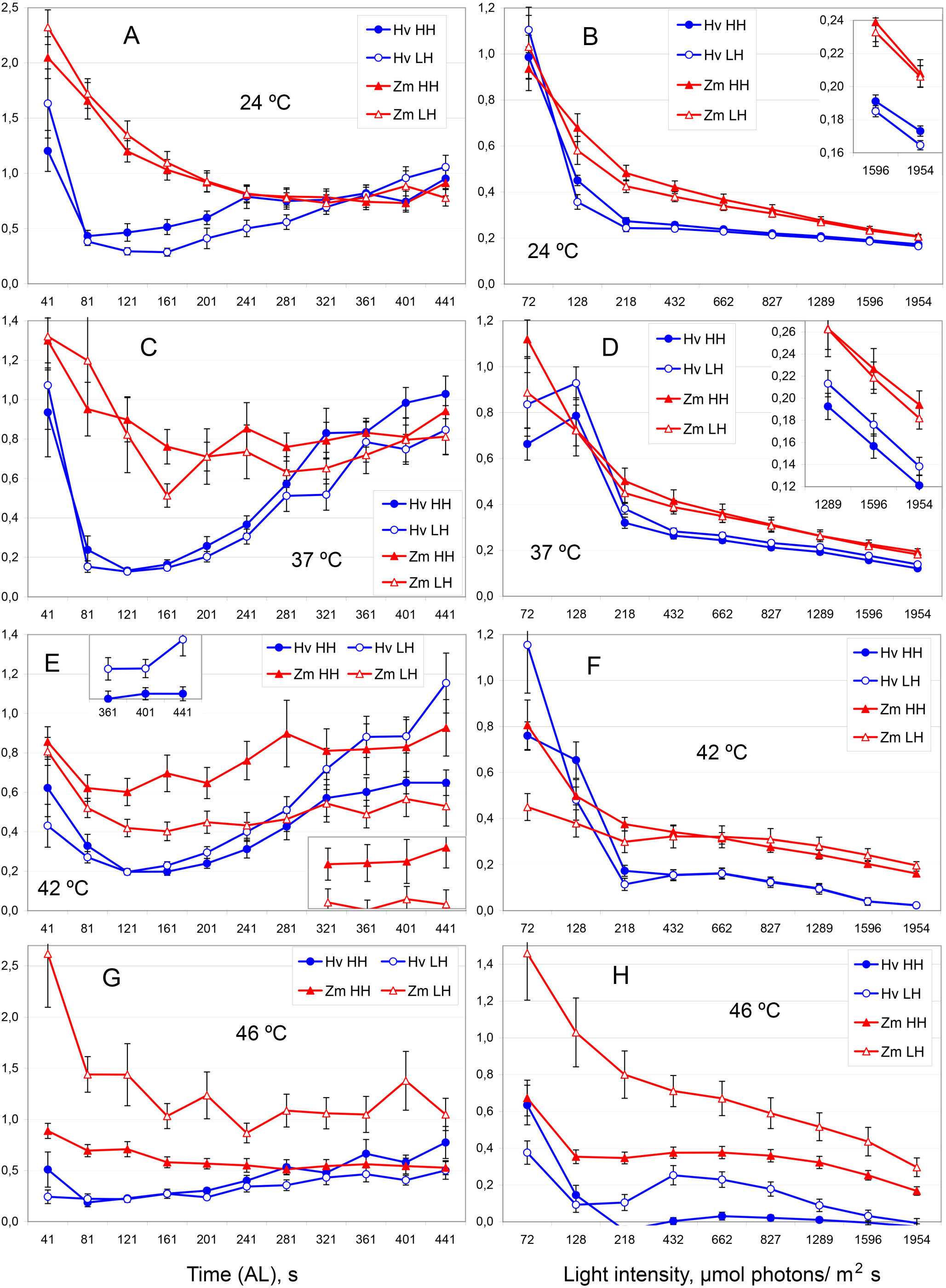
The ratio qC/Y(ND) showing balance between limitation at the acceptor sides of PSII (qC) and limitation at the donor side of PSI (Y(ND)). A, C, E, G – IC; B, D, F, H – RLC. A, B – 24°C; C, D – 37°C; E, F – 42°C; G, H – 46°C. All designations are the same as in Fig. 5. The insets show the corresponding small values with the higher resolution. All designations are the same as in Fig. 5. The same data are presented in an alternative manner for analyzing the temperature effect (Suppl. Fig. S6).

Elevating temperature stimulated a decrease of qC/Y(ND) (Suppl. Fig. S6); however, this decrease was not so obvious as the corresponding decrease in qC/Y(NA) (Suppl. Fig. S5). At 37°C, qC/Y(ND) reduced in the beginning of IC; at 42°C, qC/Y(ND) decreased in different parts of IC and RLC; at 46°C, qC/Y(ND) usually diminished, while in maize under LH conditions qC/Y(ND) ratio increased (Suppl. Fig. S6).

## 4. Discussion

### 4.1. Acceptor-side limitations of PSI and PSII

Limitations at the acceptor sides of PSII and PSI changed rather independently to each other throughout both IC and RLC with a sole exception: 1 s after AL induction, qC and Y(NA) changed in an agreement and demonstrated very similar ratios qC/Y(NA) (Fig. 1, Fig. 3, Suppl. Fig. S5). One second is a short time in slow kinetic analysis applying in the current study; in fast and ultra-fast kinetic analyses, one second is sufficient for multiple processes around PSII and PSI (Kalaji et al 2014). Figure 7 shows all the values qC, Y(NA), and qC/Y(NA) after 1 s since AL induction. We can find there three general tendencies and two species specific features.

**Fig. 7.**
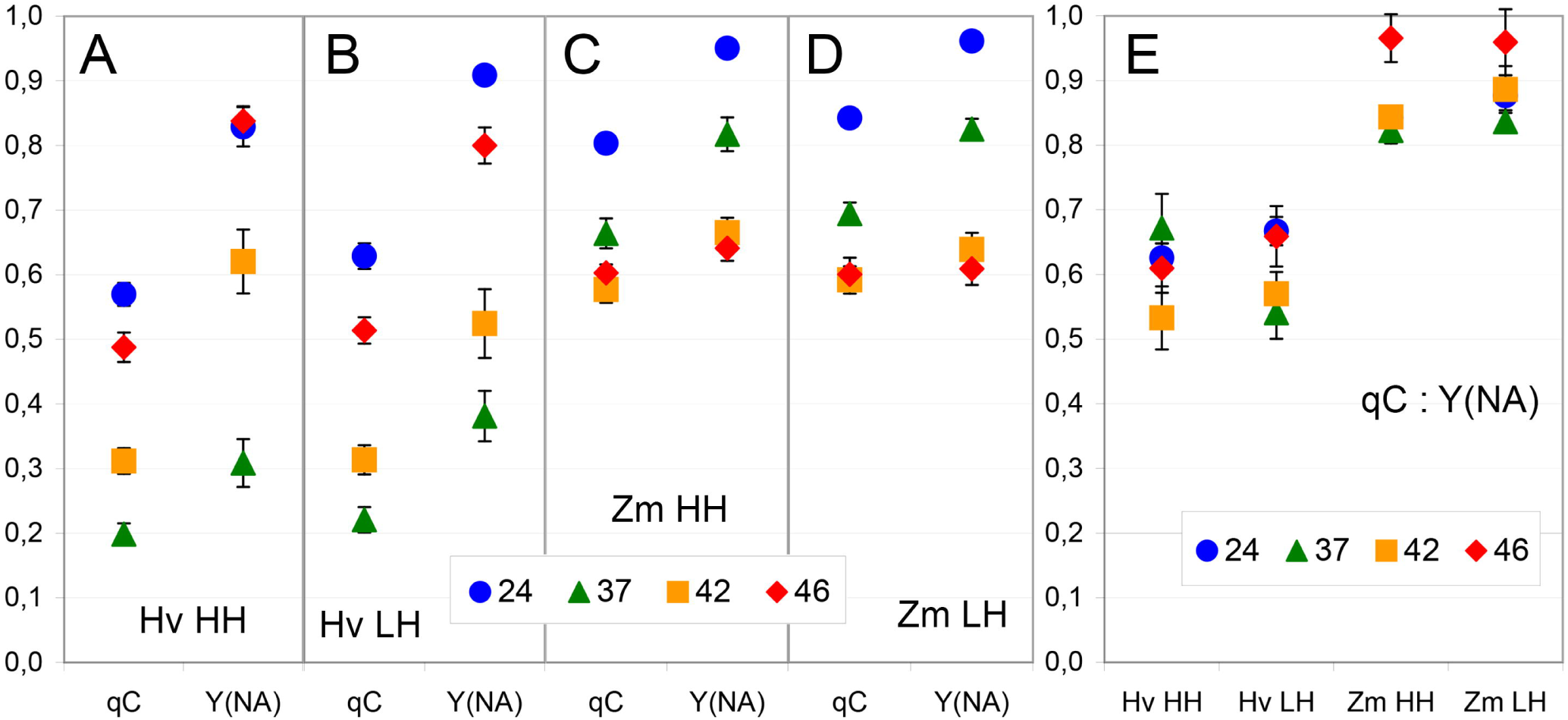
One second after AL induction: Limitations at the acceptor sides of PSII (qC) and PSI (Y(NA)). The data from Figs 1, 3, and 5. A-D – limitations at the acceptor sides of PSII (qC) and PSI (Y(NA)); E – ratio qC/Y(ND). A – barley (Hv) under HH conditions; B – barley under LH conditions, C – maize (Zm) under HH conditions; D – maize under LH conditions; E – all species and air humidity conditions. Blue circles – 24°C; green triangles – 37°C; orange squares – 42°C; red diamonds – 46°C. Means ± SE.

In both species, 1) elevating temperature decreased limitations at the acceptor sides of both PSII and PSI (Fig. 7A-D); 2) qC and Y(NA) changed with large amplitudes (Fig. 7A-D), while their ratio qC/Y(ND) varied in a limited range (Fig. 7E); 3) qC < Y(NA) observed in all but one cases (Fig. 7A-D). However, the decreasing tendency carried out in both species differently. In maize, qC and Y(NA) decreased at 37°C slightly; at the higher temperatures, the decrease enlarged (Fig. 7C,D). In barley, qC and Y(NA) decreased at 37°C greatly; at the higher temperatures, the decreasing effect exhausted and values qC and Y(NA) gradually reversed back to the control level (Fig. 7A,B). Probably, the decrease was governed by a single factor in maize while two factors governed the changes in barley: one factor decreased limitations at all elevating temperatures and another factor increased limitations starting from 42°C. The distinction between qC and Y(NA) also was different in two species. In maize, qC was slightly smaller than Y(NA); qC/Y(NA) ratios varied in the range 0.8-1 (Fig. 7C-E). In barley, qC was substantially smaller than Y(NA); qC/Y(NA) ratios varied in the range 0.5-0.7 (Fig. 7A-B,E).

The revealed agreement can be explained by a simple assumption that a single pool of molecules interacts with the acceptors sides of both PSII and PSI; this a pool is able accepting electrons, thereby reducing acceptor-side limitations of PSII and PSI for a short time. Therefore, changes of this pool influences relaxation of PSII and PSI proportionally. Oxygen, Fd, or some other molecules can be hypothesized for this role.

Oxygen accepts electrons at the acceptor sides of both PSII and PSI (Kozuleva 2022). Ferredoxin also interacts with the acceptor sides of both photosystems. At the acceptor side of PSI, Fd accepts electrons from PSI and donates them to ferredoxin-NADP reductase (FNR, EC 1.18.1.2) for NADP reduction. At the acceptor side of PSII, Fd is bound by PGR5/PGRL1 and/or NDH dehydrogenase-like complex; these complexes use Fd as an electron donor for reduction of plastoquinones (Yamori and Shikanai 2016). However, in nonphotosynthetic plastids, FNR performs the reaction in reverse direction: FNR takes an electron from NADPH to reduce Fd (reviewed in Fukuyama 2004). Possibly, PGR5/PGRL1 and/or NDH complexes are also able carrying out their reaction in the reverse direction in some exceptional cases. So, Fd accepts electrons directly from the acceptor side of PSI and may accept electrons from plastoquinones at the acceptor side of PSII in exceptional cases.

The decrease of transient acceptor-side limitations is rather specific to HS because Cd (Lysenko and Kusnetsov 2024) and other heavy metals (Lysenko, unpublished) did not impose such effect. Heat stress activated expression of Fd gene (Meng et al 2022). Increased expression of Fd genes stimulated a viability under HS and, vice versa, inhibition of Fd gene expression increased a lethality under HS (Lin et al 2015). HS inhibited FNR through nitration of tyrosine residues (Chaki et al 2011). It can be hypothesized that an increase of Fd content alleviated 1^st^-second limitations of PSI (Y(NA)) and PSII (qC) that was observed in maize at 37-46°C and in barley at 37°C (Fig. 1, Fig. 3). In barley at 42-46°C, inhibition of FNR and/or other enzymes using Fd for the reduction of sulfur and nitrogen and glutamate synthesis (Fukuyama 2004) could retain Fd in reduced state abolishing its electron accepting ability. In C_4_-plants, both HS resistance and N-metabolism are more effective (see Introduction); therefore, Fd pool could be more oxidized and ready to electron acception in maize at 42-46°C.

Excepting the peak after 1 s of AL, limitation of PSI at the acceptor side is rather small in barley, maize (Lysenko et al 2020; Lysenko and Kusnetsov 2024), and in *Arabidopsis* wild-type plants (Rantala et al 2020). This also was observed in the current study (Fig. 1). Increasing AL intensity enlarged both Y(NA) and Y(ND). At 24-42°C, Y(NA) remained minor component and Y(ND) appeared major component of PSI limitation. The impact of both Y(NA) and Y(ND) at high light (RLC) and 42°C is shown in Fig. 8A as an example; at 24°C and 37°C, the pictures are generally similar. Under nearly lethal temperature and rather high AL intensities, Y(NA) increased substantially. At 46°C and AL ≥ 432 μmol photons m^-2^ s^-1^, Y(NA) reached the value 0.4 that was similar to the value of Y(ND) (Fig. 8B). The effect was larger in maize than in barley; the effect was larger under LH than under HH conditions. Maize plants under LH conditions demonstrated even Y(NA) > Y(ND) under the highest AL intensities. Thus, under the most severe HS, processes in stroma restricted activity of the electron-transport chain greatly.

**Fig. 8.**
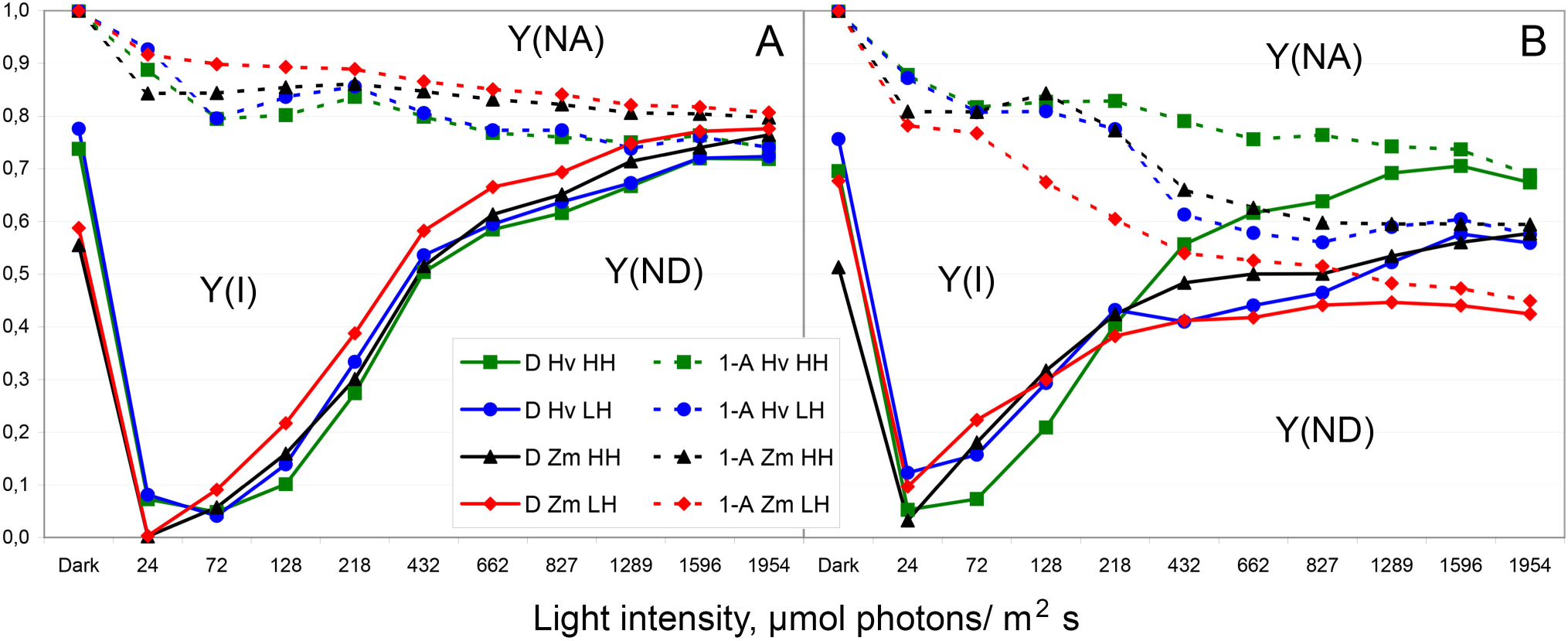
Limitations of PSI at 42°C and 46°C. The data from Figs 1 and 2. A – 42°C (pictures at 24°C, 37°C, and 42°C are generally similar); B – 46°C. Solid lines – Y(ND); the area of Y(ND) is below each line. Dashed lines – [1 – Y(NA)]; the area of Y(NA) is above each line, between a line and 1. Y(I) + Y(ND) + Y(NA) = 1 (Klughammer and Schreiber 2008); therefore, area between corresponding solid and hatched lines is 1 – Y(NA) – Y(ND) = Y(I). Green squares – barley (Hv) under HH conditions; blue circles – barley (Hv) under LH conditions; black triangles – maize (Zm) under HH conditions; red diamonds – maize (Zm) under LH conditions. Means only are shown for the sake of simplicity.

At 46°C, LH and grow light (72 μmol photons m^-2^ s^-1^) conditions, Y(ND) unambiguously increased in barley and maize in both IC (Fig. 2C,G) and RLC (Fig. 2D,H). Under the higher AL intensities in RLC, the growth of Y(ND) was restricted that coincided with the increase of Y(NA) (Fig. 1, Fig. 8). We should avoid a suggestion that high light, temperature and arid air are favorable and decreased limitation below the control level. Instead, it can be supposed that large limitation at the acceptor side of PSI relaxed someway its limitation at the donor side. The same suggestion should be applied to 1 s after AL induction when huge flash of Y(NA) relaxed Y(ND) completely (Fig. 1, Fig. 2).

The large increase of Y(NA) quasi-stationary level is rather specific to HS (Fig. 1D,F,H). Heavy metals did not cause such increase (Lysenko et al 2020; Lysenko and Kusnetsov 2024; Lysenko, unpublished); Cu-induced increase of Y(NA) was quite dissimilar (Lysenko et al 2020).

In our previous study of Cd effect, the ratio qC/Y(NA) mostly mirrored changes of qC (Lysenko and Kusnetsov 2024). In the current study, ratio qC/Y(NA) varied diversely (Fig. 5, Suppl. Fig. S5). Consequently, limitations at the acceptor sides of both photosystems changed rather independently to each other. More frequently, the values qC/Y(NA) were larger in maize than in barley (Fig. 5). Generally, HS increased Y(NA) (Fig. 1) and decreased qC (Fig. 3). Therefore, HS decreased qC/Y(NA) ratio (Suppl. Fig. S5); sometimes, the decrease was great. Probably, the increase in limitation at the acceptor side of PSI also relaxed someway limitation at the acceptor side of PSII.

### 4.2. Non-photochemical quenching

While non-photochemical quenching is a popular subject for studies, the comparison of non-photochemical quenching between C_3_- and C_4_-species remained out of focus. Recently, we demonstrated that non-photochemical quenching varied more in barley than in maize (Lysenko and Kusnetsov 2024). The current study supports this conclusion.

After AL induction, the immediate flash of acceptor-side limitation of PSII (Fig. 3B,D,F,H) was quenched by following increase of non-photochemical quenching (Fig. 4B,D,F,H). In barley, the flash of qC at 1^st^ s was followed with equal or even higher peak of qN for 40 s later (Fig. 9A-D). Both qC and qN are normalized to Fv, therefore, their values can be compared to each other; qC + X(II) + qN = 1 (Lysenko et al 2020). We showed that stress accelerates early processes in IC during readaptation to light conditions (Lysenko et al 2023; Lysenko and Kusnetsov 2024). Probably, an induction of non-photochemical quenching was slower in control than in HS-stressed plants; thereby, genuine peak of qN was achieved few seconds later (at 50-60 s) in the control and was higher. The shape of peak in control barley plants under HH conditions especially supports this suggestion (Fig. 4A) while the corresponding peak under LH conditions (Fig. 4C) does not contradict to this assumption. Maize plants induced rather small increase of qN at 41^st^ s; this increase at 41^st^ s did not represent a peak and was rather equal in all the variants despite the changes of qC at 1^st^ s in maize plants (Fig.9E-F).

**Fig. 9.**
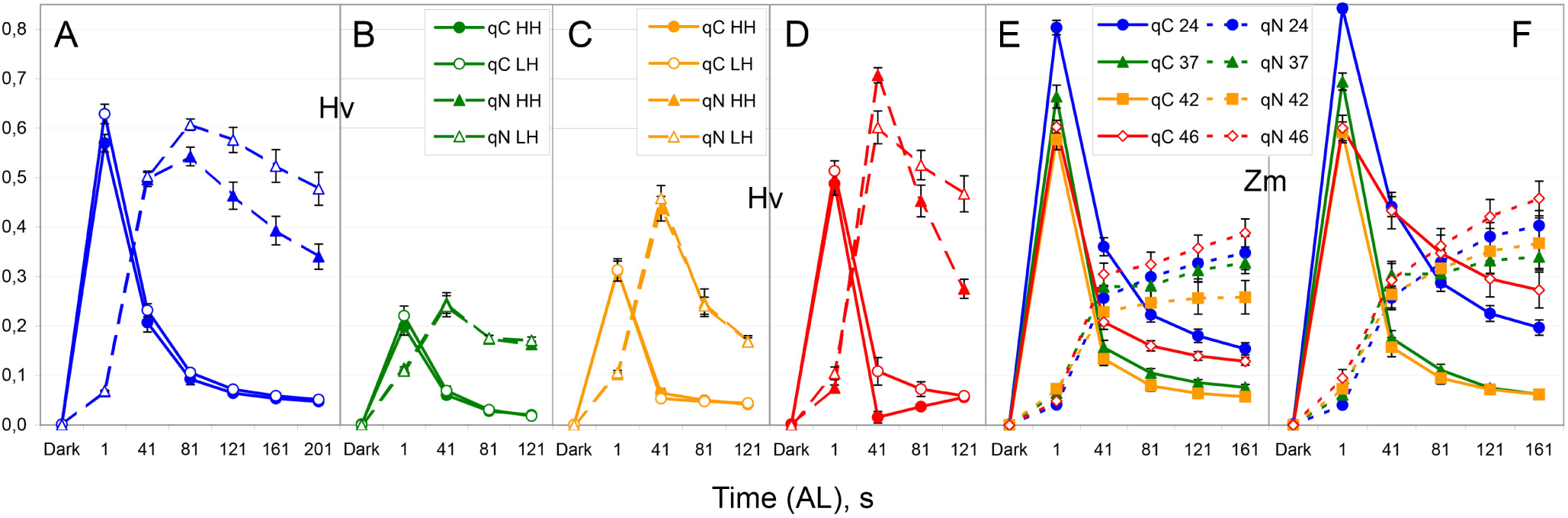
Donor-side limitation (qC) and non-photochemical quenching (qN) of PSII in the beginning of IC. The data from Figs 3 and 4. A-D – barley (Hv). A – 24°C; B – 37°C; C – 42°C; D – 46°C. Solid lines with circles – qC; dashed lines with triangles – qN; filed symbols – HH conditions; open symbols – LH conditions. E – maize (Zm) under HH conditions; F – maize (Zm) under LH conditions. Solid lines – qC; dashed lines – qN; filled blue circles – 24°C; filled green triangles – 37°C; filled orange squares – 42°C; open red diamonds – 46°C. Means ± SE.

So, barley plants induced regulated flash of non-photochemical quenching corresponding in size to the previous flash of limitation at the donor side of PSII (Fig. 9A-D), while maize plants increased non-photochemical quenching in an unregulated manner to a constant value (∼ 0.25-0.3) irrespectively to sizes of the previous flashes of limitation at the donor side of PSII (Fig. 9A-D).

When IC dynamics reached the steady state levels, a similar effect was observed in both species. At 46°C, LH conditions similarly decreased the photochemical quenching of PSII in barley and maize (Lysenko et al 2023). However, this downregulation was underlain with different mechanisms. At 46°C in LH conditions, barley plants increased the steady state level of non-photochemical quenching qN (Suppl. Fig. S4G), while maize plants increased the steady state level of closed PSII qC (Suppl. Fig. S3G). In maize, a faint increase of qN was observed under LH conditions (Suppl. Fig. S4G); the pairwise t-test comparison showed no differences, while whole stationary dynamics (81-441 s) were significantly different according to two-tail paired nonparametric binomial test (p<0.01). So, non-photochemical component was very small in maize.

On the whole, non-photochemical quenching varied to a larger extent in barley than in maize (Fig. 3). Under tolerated HS (37-42°C), both plant species generally decreased non-photochemical quenching (Fig. 3), probably, to minimize an additional warming through the thermal dissipation of light energy that is major component of non-photochemical quenching. Under nearly lethal HS (46°C), qN mostly increased (Fig. 3); possibly, severe stress exhausted plants’ ability to minimize thermal dissipation.

### 4.3. Limitations between PSII and PSI

The fragment of electron transport chain between PSII and PSI consists of plastoquinones, cytochrome b_6_f complex, and plastocyanin. A priori, we assume that this fragment should impose same or similar limitations at both ends: at the acceptor side of PSII (qC) and at the donor side of PSI (Y(ND)). Hence, qC and Y(ND) should change proportionally to each other while their ratio qC/Y(ND) should be rather stable.

Recently, we demonstrated that qC/Y(ND) ratio has its own dynamics in both IC and RLC. In the beginning of both IC and RLC, the ratio qC/Y(ND) decreased then stabilized for a short time (quasi-stationary level); next, constant AL intensity in IC enables increasing qC/Y(ND) while increasing AL intensity in RLC causes decreasing qC/Y(ND) (Lysenko and Kusnetsov 2024). Quasi-stationary fragments of qC/Y(ND) dynamics reflect proportional changes of qC and Y(ND); in most cases, a single variant demonstrated quasi-stationary levels at same values of qC/Y(ND) in both IC and RLC (Lysenko and Kusnetsov 2024). The ratios qC/Y(ND) were calculated in the current study; they altered our initial conclusions.

Dynamics corresponding to an intuitive assumption of rather stable qC/Y(ND) were found in IC. After AL induction, several qC/Y(ND) curves demonstrated small changes in the beginning and rather horizontal tendency later; such nearly plane curves were observed mostly in maize at 37-46 °C (except, 46°C LH) and in barley at 46°C and LH conditions (Fig. 6C,E,G). Previously described IC pattern “decrease – short plateau – increase” was observed in all barley variants excepting at 46°C and LH conditions (Fig. 6A,C,E,G); the later stage of increase frequently was much larger than it was observed previously (Lysenko and Kusnetsov 2024). Maize in control and at 46°C and LH conditions demonstrated large decrease and plateau with no increase later (Fig. 6A,G).

RLC dynamics (Fig. 6B,D,F,H) mostly corresponded to the previously described pattern “decrease – short plateau – decrease” (Lysenko and Kusnetsov 2024). Frequently, “plateau” was absent and dynamics looked like “fast decrease – slow decrease” (e.g., maize at 24-37°C, see Fig. 6B,D). In some cases, “plateau” tended to be transfomed to “wave” (clear case barley at 46°C and LH conditions, Fig. 6H)

Usually, short quasi-stationary levels were at different levels in IC and RLC (Fig. 6). Rarely, both IC and RLC “plateaus” were at the same level in a single variant; it was observed in barley, mostly under LH conditions (e.g., barley at 42°C, Fig. 6E,F). Though, short quasi-stationary levels overlapped in dynamics. In IC, quasi-stationary levels were at (81-)121-161 s (“bottoms”) in barley and at 281-321-361 s (“horizon”) in maize. In RLC, control barley plants demonstrated quasi-stationary levels at 218-432 μmol photons m^-2^ s^-1^; at 42-46°C, quasi-stationary levels were observed at 432-662 μmol photons m^-2^ s^-1^ in both species (Fig. 6B,D,F,H).

Why qC/Y(ND) curves obtained previously (Lysenko and Kusnetsov 2024) and in the current study (Fig. 6) are so different? Partially, the controversy could be caused by the different factors studied – Cd and HS. Controls in both studies have a limited similarity. However, more similarity to previously obtained data (Lysenko and Kusnetsov 2024) showed RLC measured at 42°C (Fig. 6F). Three more differences should be critical. 1) In the previous study (Cd), plants were grown in phytotron; in the current work (HS), plants were grown in a limited volume of thermostat chambers that can influence gas exchange with both CO_2_ and signaling molecules. 2) Plants were adapted to different grow light – 180-220 μmol photons m^-2^ s^-1^ (Cd) and 60-80 μmol photons m^-2^ s^-1^ (HS); therefore, IC were measured at different AL intensity – 128 μmol photons m^-2^ s^-1^ (Cd) and 72 μmol photons m^-2^ s^-1^ (HS). 3) Different sets of AL intensities were used in RLC: [24]/(39)-72-(97)-128-(168)-218-(341)-432-662-827-(1030)/[1289]-[1596]-1954 – where AL intensities solely used in the previous work (Cd) marked with round parenthesis and intensities solely used in the current work marked with rectangular parenthesis. Until 432 μmol photons m^-2^ s^-1^, AL increase was twice more gradual in the previous (Cd) then in the current work (HS). Probably, more longer and/or gradual increase of AL intensities is preferable for revealing inflection points in qC/Y(ND) dynamics in RLC.

The previous and the current works demonstrated that limitations between PSII and PSI can vary diversely. The existing data set is insufficient for conclusion making; fortunately, some of the previous conclusions were supported in this study. It is worth to pay more attention to the analysis of qC/Y(ND) dynamics in further studies.

## Supporting information

Supplemental Tabs & Figs

## Acknowledgement

The author thanks Dr. Klaus A.A. for growing the plants and Dr. Kozuleva M.A for caring out the measurement by Dual-PAM-100 in 3 out of 4-5 biological experiments. The seeds of maize were kindly provided by Dr. E.V. Kartamysheva (VNIIMK, Don experimental station). The research was performed within the state assignment of Ministry of Science and Higher Education of the Russian Federation (theme No. 122042700044-6).

## Abbreviations

AL: actinic light
BS: bundle sheath (cells or chloroplasts)
Chl: chlorophyll
Fd: ferredoxin
HH: higher (relative) humidity (of air)
HS: heat stress
IC: induction curve
LH: lower (relative) humidity (of air)
M: mesophyll
NAD(P)-ME: NAD(P)-dependent malic enzyme
PAM: pulse amplitude modulation
PEP-CK: phosphoenolpyruvate carboxykinase
PSI, PSII: photosystem I and II
P_700_: reaction center Chl of PSI
RLC: rapid light curve
SE: standard error
SP: saturation pulse

