## Supplemental Tabs & Figs for "Differences between barley and maize revealed in limitations of photosystems I and II under high temperature and low air humidity"

by *Eugene A. Lysenko*

**Source:** bioRxiv

### General map of deleted points for calculating the ratios qC/Y(ND) and qC/Y(NA).

**Table S1**. The ratio qC/Y(ND) in maize. Map of points deleted because of small denominator value.

| | 24ºC | | 37ºC | | 42ºC | | 46ºC | |
|---|---|---|---|---|---|---|---|---|
| | HH | LH | HH | LH | HH | LH | HH | LH |
| Whole IC deleted: | | | 3 | 2 | 4 | 2 | 1 | |
| Number of IC: | 24 | 23 | 17 | 18 | 15 | 14 | 17 | 16 |
| 41 | | | 1 | | | | | 1 |
| 81 | | | **1** | | | | | |
| 121 | | | | **1** | | | | |
| 161 | | | **1** | **1** + 1 | | | | |
| 201 | | | | | | | | |
| 241 | **1** | | | | | | | 1 |
| 281 | **1** | | | **1** | **2** | | | |
| 321 | | 1 | 1 | **1** | **1** | 1 | | |
| 361 | **2** | | | | **3** | **1** | | 1 |
| 401 | **2** | | **1** | **2** | **3** | 1 | | |
| 441 | 7 * + **1** | 7 * + 1 | **1** | 1 | **2** | 1 + **3** | | **1** |
| Whole RLC deleted: | | | | | | | | |
| Number of RLC: | 23 | 23 | 20 | 20 | 19 | 16 | 18 | 16 |
| 72 | **3** | 1 | 2 + **2** | **3** | 2 + **3** | **3** | 1 | 1 |
| 128 | | | 1 | | | | 1 | |
| 218 | | | | | | | | |
| 432 | | | | | | | | |
| 662 | | | | | | | | |
| 827 | | | | | | | | |
| 1289 | | | | | | | | |
| 1596 | | | | | | | | |
| 1954 | | | | | | | | 1 * |

Deleted Y(ND): negative value; zero value (0);
**value ≤ 0.025**; **0.025 < value > 0.030**; value ≥ 0.030.
* - data absent.

Table S2. The ratio qC/Y(ND) in barley. Map of points deleted because of small denominator value.

| | 24ºC | | 37ºC | | 42ºC | | 46ºC | |
|---|---|---|---|---|---|---|---|---|
| | HH | LH | HH | LH | HH | LH | HH | LH |
| Whole IC deleted: | 6 | 2 | 3 | 3 | 1 | | | |
| Number of IC: | 19 | 24 | 16 | 18 | 17 | 19 | 17 | 22 |
| 41 | 1 + **1** | 2+**1**+1+1 | **2** | **1**+1+1 | | 1 * | 9 + 1 | 2+2+**1**+2 |
| 81 | | | **1** | **2** + 2 | | 1 * | | 2 + **1** + 2 |
| 121 | | | | | | | | |
| 161 | | 2 | | | | | | |
| 201 | **1** | **2** | | | | | 1 + 1 | 1 |
| 241 | | | | **1** | | | **1** | |
| 281 | **2** | | **2** | 1 + **1** | | | 1 | |
| 321 | **3** | **1** | 1 + **3** | 1 + **6** | | | | |
| 361 | 1 + **1** | 1 + **3** | 1 + **2** | 1 + **7** | **1** + 1 | **1** | | **1** |
| 401 | 2 + **5** | 1 + **3** | 2 + **6** | 1 + **8** | **1** + 1 | **1** | 1 + **1** | 1 + **1** |
| 441 | 5 * + **7** | 5 * + **6** | 1 + **6** | 3 + **9** | **3** | 1 + **2** | 2 + **1** | **1** |
| Whole RLC deleted: | | | | | | | | |
| Number of RLC: | 25 | 25 | 19 | 21 | 17 | 19 | 17 | 20 |
| 72 | 2 + **8** | 1 + **5** | 4+**7**+1 | 4 + **10** | **3** + 1 | 2 + **6** | 2 + **1** | 2 |
| 128 | | | **2** | | | | | |
| 218 | | | | | | | | |
| 432 | | | | | | | | |
| 662 | | | | | | | | |
| 827 | | | | | | | 1 * | |
| 1289 | | | | | | | 1 * | |
| 1596 | | | | | | | 1 * | |
| 1954 | | | | | | | 1 * | |

Deleted Y(ND): negative value; zero value (0);
**value ≤ 0.025**; **0.025 < value > 0.030**; value ≥ 0.030.
* - data absent.

Table S3. The ratio qC/Y(NA) in maize. Map of points deleted because of small denominator value.

| | 24ºC | | 37ºC | | 42ºC | | 46ºC | |
|---|---|---|---|---|---|---|---|---|
| | HH | LH | HH | LH | HH | LH | HH | LH |
| Whole IC deleted: | | | 1 | 1 | | 3 | 1 | 2 |
| Number of IC: | 24 | 23 +1# | 19 | 19 | 19 | 14 | 17 | 14 |
| 1 | | | | | | | | 1 * |
| 41 | | | | | **2** | | | |
| 81 | | | | | 1 + **3** | | 1 + **3** | |
| 121 | | | | | 1 + **1** | **1** | | |
| 161 | | | | 1 | 1 + **1** | **1** | 1 | 1 + 1 |
| 201 | | | | | **1** | 1 | 1 | **1** |
| 241 | | | | **1** | 1 + **1** | | | |
| 281 | | | | | | **1** | | 1 |
| 321 | | | | **1** | | | 2 | |
| 361 | | | | **1** | | | 1 | |
| 401 | | | | | | | | |
| 441 | 7 * | 7 * | | | **1** | | | |
| Whole RLC deleted: | | | | 1 | | 4 | | |
| Number of RLC: | 23 | 23 | 20 | 19 | 19 | 12 | 18 | 16 |
| 72 | | | 1 | **1** | | | 1 | 1 |
| 128 | | | | | | | 1 + **1** | 1 |
| 218 | | | | | | | 1 | |
| 432 | | | **1** | | **2** | **1** | 1 | |
| 662 | | | **1** | | | | | 1 |
| 827 | | | **1** | | | | | 1 |
| 1289 | | | 1 | | | | | |
| 1596 | | | **1** | | | | | |
| 1954 | | | | | | | | 1 * |

Deleted Y(NA): negative value; zero value (0);
**value ≤ 0.025**; **0.025 < value > 0.030**; value ≥ 0.030.
* - data absent. # - incomplete (short) curve.

Table S4. The ratio qC/Y(NA) in barley. Map of points deleted because of small denominator value.

| | 24ºC | | 37ºC | | 42ºC | | 46ºC | |
|---|---|---|---|---|---|---|---|---|
| | HH | LH | HH | LH | HH | LH | HH | LH |
| Whole IC deleted: | 2 | 2 | | 1 | | 1 | 1 | 4 |
| Number of IC: | 23 | 24 | 19 | 20 | 18 | 18 | 16 | 18 |
| 1 | 1 * | 1 * | | | | 2 * | 1 * | |
| 41 | | | 1 | | | 1 * | 8+1+1 | 2 + 1 |
| 81 | 1 | 1 | 2 | 3 | | 1 * | | 2 |
| 121 | 1 | 1 + 3 | 1+1+1 | 1 | 1 | 1 | 1 + 1 | |
| 161 | 1 | 1 + 2 | 1 + | 1 | | | | 1 + 1 |
| 201 | 1 | 1 + 1 | | 2 | | | 2 | 1 |
| 241 | 1 | 2 | 1 + | 1 + 1 | | | 2 | 1 |
| 281 | | 1 + 1 | | 1 | | | 2 | |
| 321 | | 1 | | 1 | | | | |
| 361 | | | | 1 | | | 1 | |
| 401 | 1 | | | | | | | |
| 441 | 7 * | 6 * + 1 | | 1 | | | | |
| Whole RLC deleted: | 2 | 2 | | 1 | | | | |
| Number of RLC: | 23 | 23 | 19 | 20 | 17 | 19 | 17 | 20 |
| 72 | | | | | | | 1 | 3 |
| 128 | 1 + 3 | 2 | | | | | 1 | 3 |
| 218 | 1 | 1 + 2 | 4 | 1 | | | | 1 + 1 |
| 432 | 1 + 1 | | 2 | 1 | | | | |
| 662 | 1 | | 1 | 1 | | 1 | | |
| 827 | | | | | | | 1 * | |
| 1289 | | | | | | | 1 * | |
| 1596 | 1 | | | | | | 1 * | |
| 1954 | | | | 1 | | | 1 * | |

Deleted Y(NA): negative value; zero value (0);
**value ≤ 0.025**; **0.025 < value > 0.030**; value ≥ 0.030.
* - data absent.

Fig. S1. Limitation at the acceptor side of PSI (Y(NA)). Revisualization of the data from Fig.1.

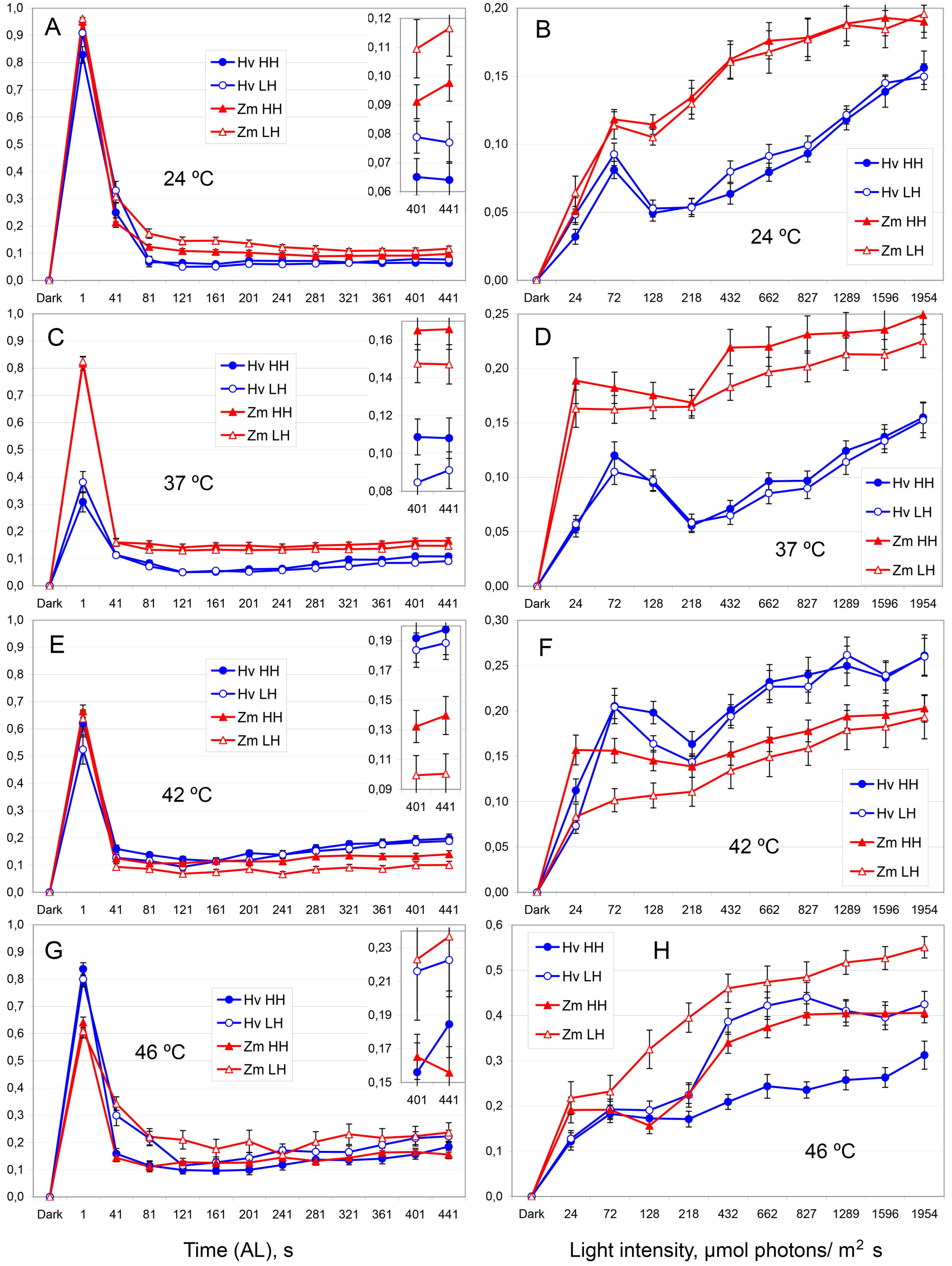


Left column – IC; right column – RLC. The temperatures are shown on panels. Blue circles – barley (Hv); red triangles – maize (Zm); filled symbols – HH air; open symbols – LH air. Means ± SE. Insets show the corresponding small values with the higher resolution.

Fig. S2. Limitation at the donor side of PSI (Y(ND)). Revisualization of the data from Fig. 2.

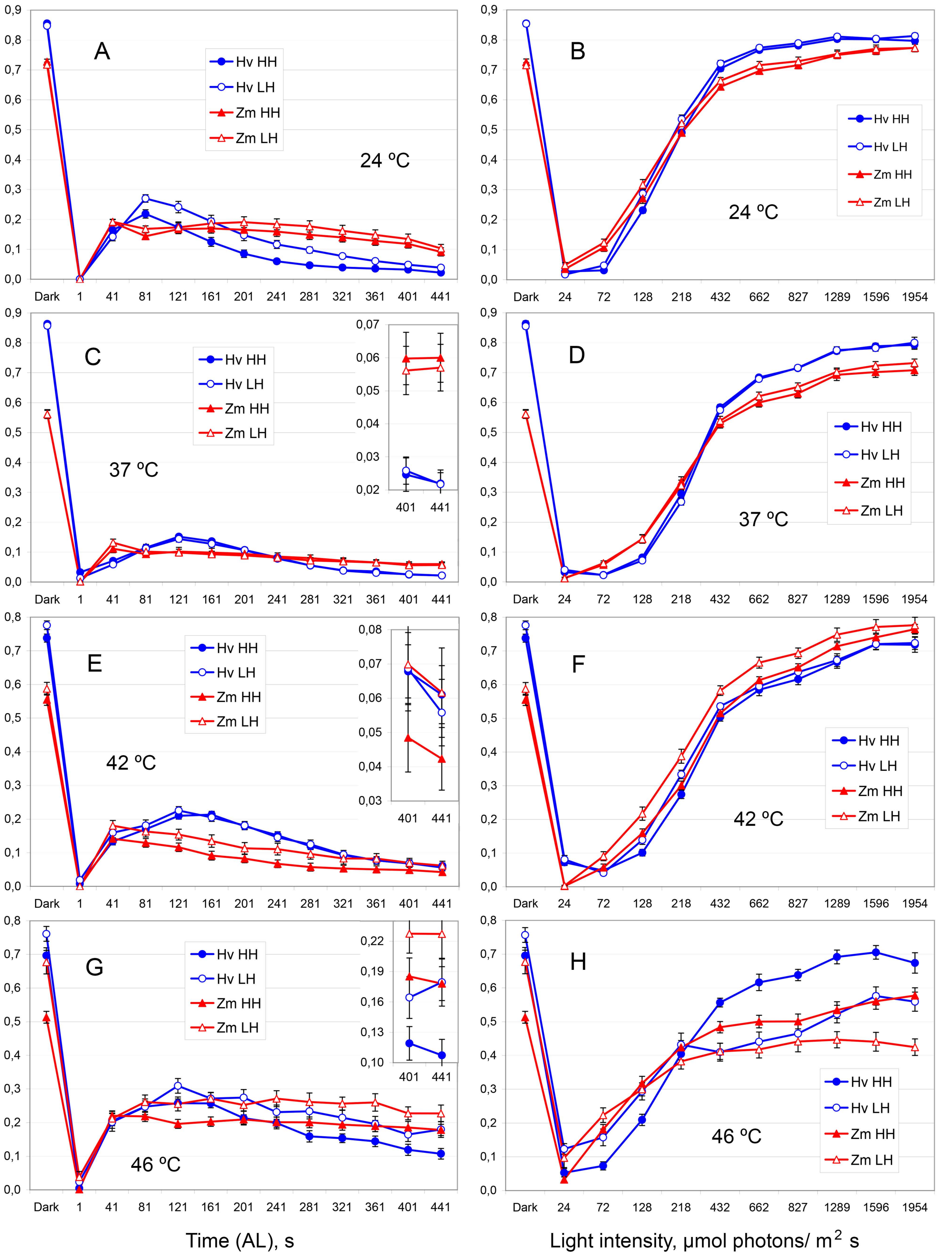


Left column – IC; right column – RLC. The temperatures are shown on panels. All designations are the same as in Fig. S1 and Fig. 5. Means ± SE. Insets show the corresponding small values with the higher resolution.

Fig. S3. Limitation at the acceptor side of PSII (qC). Revisualization of the data from Fig. 3.

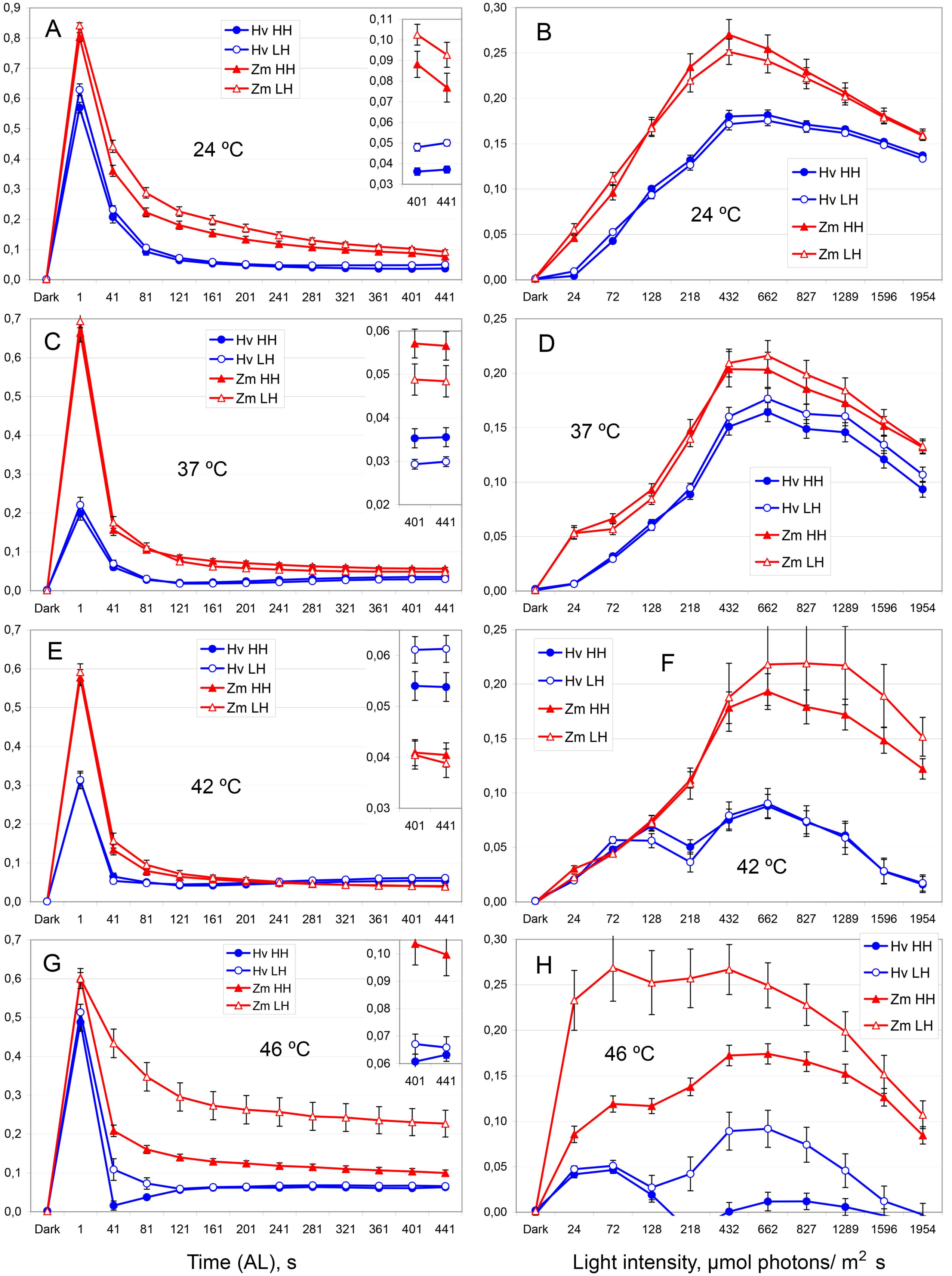


Left column – IC; right column – RLC. The temperatures are shown on panels. All designations are the same as in Fig. S1 and Fig. 5. Means ± SE. Insets show the corresponding small values with the higher resolution.

Fig. S4. Non-photochemical quenching of PSII (qN). Revisualization of the data from Fig. 4.

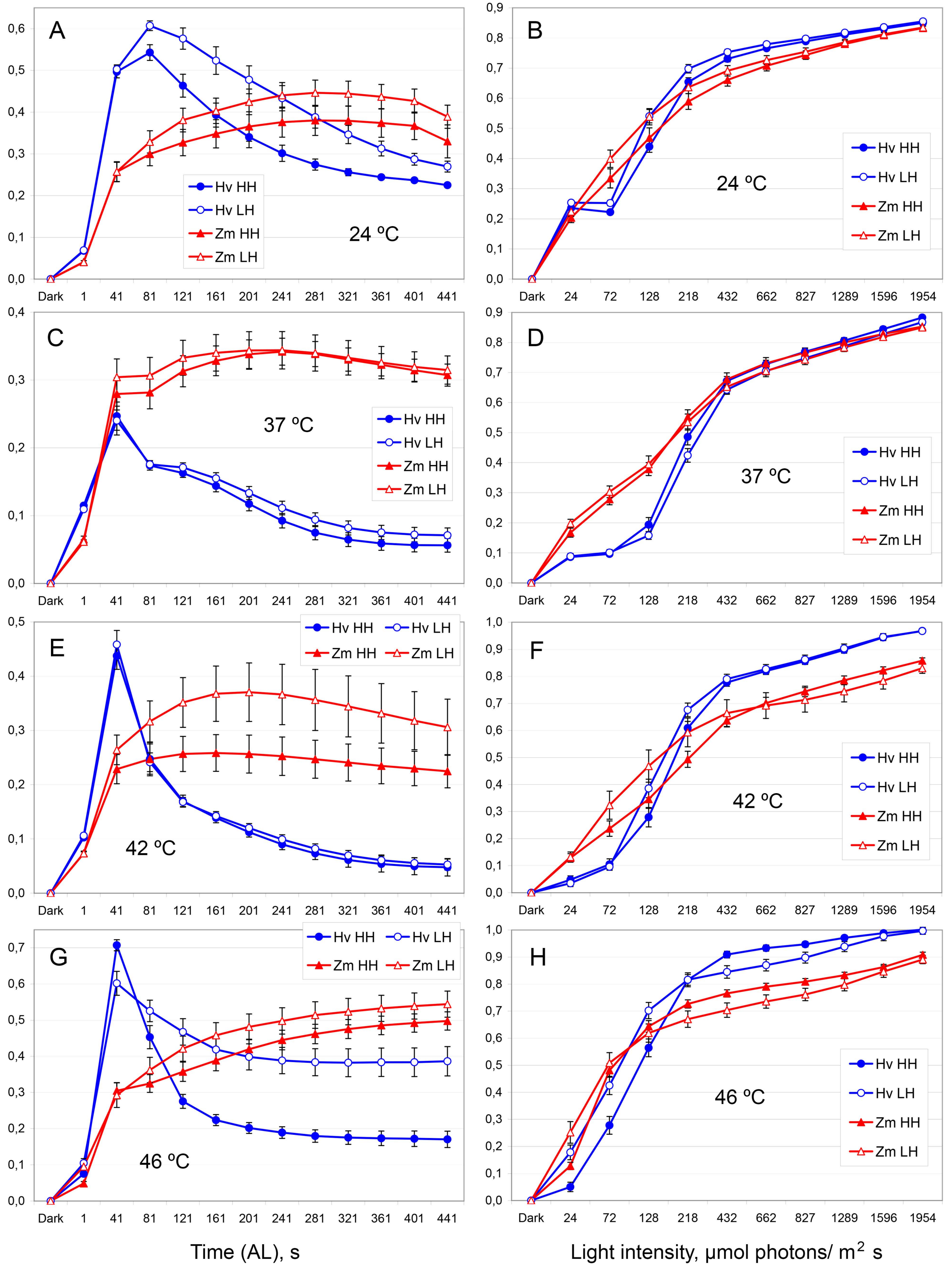


Left column – IC; right column – RLC. The temperatures are shown on panels. All designations are the same as in Fig. S1 and Fig. 5. Means ± SE.

Fig. S5. The ratio qC/Y(NA). Revisualization of the data from Fig. 5.

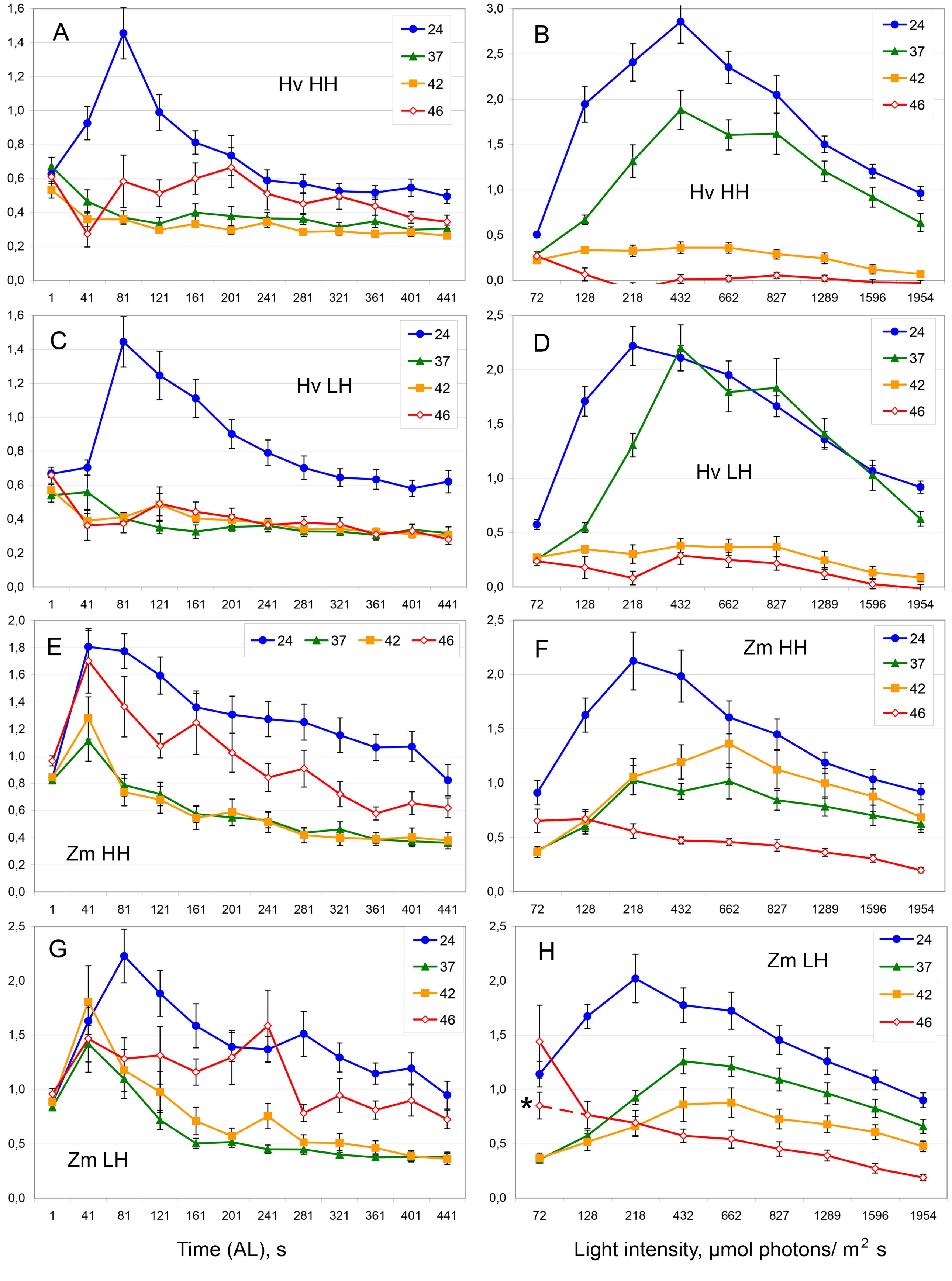


Left column – IC; right column – RLC. The variants are shown on panels: Hv – barley; Zm – maize; HH and LH – higher and lower relative humidity of air. Numerals indicate temperature in Celsius degree. * – variant of corresponding point in curve without 3 drastically different but irremovable data points (as more credible; connected with dashed line). Means ± SE.

Fig. S6. The ratio qC/Y(ND). Revisualization of the data from Fig. 6.

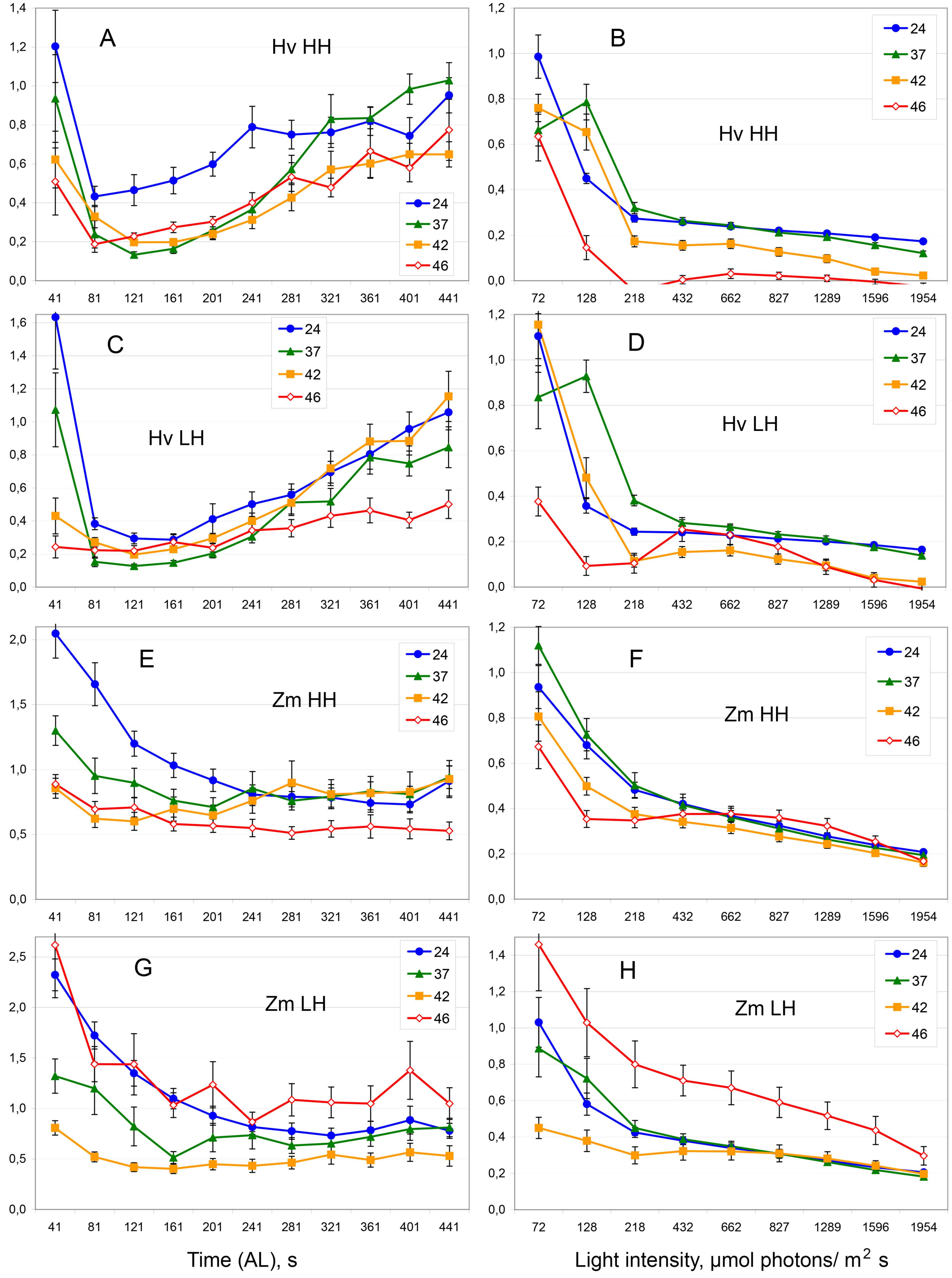


Left column – IC; right column – RLC. The variants are shown on panels. All designations are the same as in Fig. S5 and Fig. 1. Means ± SE.
